# Cocaine-induced DNA-PK relieves RNAP II pausing by promoting TRIM28 phosphorylation

**DOI:** 10.1101/2024.08.19.608673

**Authors:** Adhikarimayum Lakhikumar Sharma, Priya Tyagi, Meenata Khumallambam, Mudit Tyagi

## Abstract

Drug abuse continues to pose a significant challenge in HIV control efforts. In our investigation, we discovered that cocaine not only upregulates the expression of DNA-dependent protein kinase (DNA-PK) but also augments DNA-PK activation by enhancing its phosphorylation at S2056. Moreover, DNA-PK phosphorylation triggers the translocation of DNA-PK into the nucleus. The finding that cocaine promotes nuclear translocation of DNA-PK further validates our observation of enhanced DNA-PK recruitment at the HIV long terminal repeat (LTR) following cocaine exposure. By activating and facilitating the nuclear translocation of DNA-PK, cocaine effectively orchestrates multiple stages of HIV transcription, thereby promoting HIV replication. Additionally, our study indicates that cocaine-induced DNA-PK promotes hyper-phosphorylation of RNA polymerase II (RNAP II) carboxyl-terminal domain (CTD) at Ser5 and Ser2 sites, enhancing both initiation and elongation phases, respectively, of HIV transcription. Cocaine’s enhancement of transcription initiation and elongation is further supported by its activation of cyclin-dependent kinase 7 (CDK7) and subsequent phosphorylation of CDK9, thereby promoting positive transcriptional elongation factor b (P-TEFb) activity. We demonstrate for the first time that cocaine, through DNA-PK activation, promotes the specific phosphorylation of TRIM28 at Serine 824 (p-TRIM28, S824). This modification converts TRIM28 from a transcriptional inhibitor to a transactivator for HIV transcription. Additionally, we observe that phosphorylation of TRIM28 (p-TRIM28, S824) promotes the transition from the pausing phase to the elongation phase of HIV transcription, thereby facilitating the production of full-length HIV genomic transcripts. This finding corroborates the observed enhanced RNAP II CTD phosphorylation at Ser2, a marker of transcriptional elongation, following cocaine exposure. Accordingly, upon cocaine treatment, we observed elevated recruitment of p-TRIM28-(S824) at the HIV LTR. Overall, our results have unraveled the intricate molecular mechanisms underlying cocaine-induced HIV transcription and gene expression. These findings hold promise for the development of highly targeted therapeutics aimed at mitigating the detrimental effects of cocaine in individuals living with HIV.

**Highlights of the study:**

a. Cocaine upregulates both the expression and activity of DNA-PK.
b. Cocaine augments the phosphorylation of DNA-PK selectively at S2056, a post-translational modification that marks functionally active form of DNA-PK.
c. Cocaine enhances the nuclear translocation of DNA-PK.
d. The DNA-PK inhibition severely impairs HIV transcription, replication, and latency reactivation.
e. Cocaine facilitates the initiation and elongation phases of HIV by enhancing RNAPII CTD phosphorylation at Ser5 and Ser2, respectively, by stimulating DNA-PK.
f. Cocaine also supports initiation and elongation phases of HIV transcription by stimulating CDK7 (the kinase of TFIIH) and CDK9 (the kinase subunit of P-TEFb), respectively.
g. Cocaine-mediated activation of DNA-PK relieves RNAP II pausing by reversing the inhibitory effect of pausing factor TRIM28 and converting it into a transactivator by catalyzing its phosphorylation at S824 site.
h. Thus, cocaine, by activating DNA-PK, facilitates the multiple phases of HIV transcription, namely, initiation, RNAP II pause-release, and elongation.

## 1. Introduction

The onset of acquired immunodeficiency syndrome (AIDS), triggered by Human Immunodeficiency Virus type 1 (HIV), is one of the most profoundly impactful diseases humanity has faced. Since the identification of HIV in 1981, extensive endeavors have been undertaken to combat HIV infection. These efforts have catalyzed significant progress in the realms of immunology and HIV virology, marking notable advancements along the way [1–4]. However, HIV eradication or a preventive vaccine is yet to be developed [5]. The current anti-HIV drug regimens (anti-retroviral therapy, ART) have been highly successful in lowering HIV/AIDS-related mortality and improving the quality of life for people living with HIV (PLWH) [1, 6]. As ART can effectively diminish the viral load to undetectable levels through standard methodologies, the substantially decreased levels of HIV while on ART facilitates the restoration and sustenance of a robust immune system. This restoration enables the body to effectively defend against opportunistic infections and illnesses [7–9]. In addition, ART greatly reduces the risk of HIV transmission [7]. On other hand, dangerous behavior, such as unprotected sex and needle sharing by illicit drug users, significantly increases HIV transmission risk [10, 11]. Although there has been remarkable achievement in controlling HIV, the prevalence of illicit drug usage remains a significant contributor to new HIV infection due to their perilous behavior [12–17]. Cocaine (Coc), a powerfully addictive stimulant drug has a high potential for abusing tendency [18–21]. Cocaine is primarily used orally, intranasal, intravenously, or by inhalation [22]. Continuous use of cocaine interferes with normal brain function; thus, it compromises judgment and decision-making capability, leading to risky behavior such as needle sharing and sexual behavior, including trading sex for drugs [23, 24]. Once infected, cocaine further increases the severity of the HIV infection; stimulates HIV replication, including in the central nervous system (CNS); and accelerates the occurrence of neurocognitive impairments [25–28]. Studies have also documented that cocaine use accelerates CD4+ T cell loss, even in ART-treated individuals [29, 30]. However, the precise mechanisms by which cocaine and HIV synergize to compromise the health of individuals living with HIV (PLWH) remain unclear.

Similar to host cell gene transcription, RNA polymerase II (RNAP II) is required for HIV transcription. RNAP II is regulated by specific phosphorylation events in the carboxyl-terminal domain (CTD) of RNAP II large subunit [31]. The human RNAP II CTD consists of 52 tandem repeats of a consensus sequence Tyr1-Ser2-Pro3-Thr4-Ser5-Pro6-Ser7 [32–35]. Many known kinases can phosphorylate RNAP II CTD. However, most notable kinases that phosphorylates RNAP II are cyclin-dependent kinase 7 (CDK7) that phosphorylate RNAP II at Ser5 and CDK9 that phosphorylate RNAP II at Ser2 [31, 34, 36]. Our previous studies documented that DNA-PK can phosphorylate RNAP II CTD in all three serine residues (Ser2, Ser5, and Ser7) [37]. We have also shown that transactivator of transcription (Tat) protein, which is vital for HIV transcription, is a potential substrate of DNA-PK [37]. Data generated from our previous study also suggested that cellular activation augments both the nuclear translocation and HIV LTR recruitment of DNA-PK [37–39].

DNA-PK, a protein kinase, requires association with DNA to become catalytically active. DNA-PK holoenzyme consisting of two components: a 450 kDa catalytic subunit (DNA-PKcs), which is a serine/threonine kinase, and a regulatory component known as Ku, a heterodimer of Ku70 and Ku80 [40–42]. DNA-PK is well studied for its role in repairing DNA damage and maintaining the stability of the genome, including during V(D)J recombination [43–45]. DNA-PK especially plays a crucial role in the non-homologous end joining (NHEJ) DNA repair pathway [46]. While multiple recent studies, including our own, have suggested a potential involvement of DNA-PK in transcriptional regulation [37, 47], the precise role of DNA-PK in the transcription process was delineated by our research [37]. It has been documented that DNA-PK interacts with various transcription factors and components of the transcription machinery [47]. Notably, DNA-PK not only engages with numerous transcription factors, such as TFIIH, P-TEFb, p53, NF-κB, and SP1, but also modulates their activity through phosphorylation. These interactions typically amplify the expression of genes regulated by these transcription factors.

During HIV transcription, phosphorylation of RNAP II CTD at the position Ser5 is associated with the early stages of transcription, particularly transcription initiation. This modification recruits capping enzyme complexes that add a 7-methylguanosine cap to the nascent RNA molecule, which protects RNA from degradation and later facilitates its processing. However, phosphorylation of RNAP II CTD at Ser2 is linked to the elongation phase of transcription, as this post-translational modification of RNAP II makes it processive or elongation-proficient, as it reduces the slipping of RNAP II from DNA template. This modification also facilitates the recruitment of transcription factors involved in mRNA maturation and processing, including splicing and polyadenylation. For efficient transcription elongation, not only processive RNAP II is required, but also the removal of negative transcription factors (NFs) that promote promoter-proximal pausing of RNAP II is essential [48–52]. Analogous to cellular gene expression, HIV Transcriptional initiation also halts after generating short nascent mRNA of around 60 nucleotides due to the binding of negative transcription factors (NFs) at the HIV LTR [53–55]. Some notable NFs are the negative elongation factor (NELF) and the 5,6-dicholoro-1-β-d-ribofuranoxylbenzimidazole (DRB) sensitivity-inducing factor (DSIF) [56, 57]. Recently, in addition to DSIF and NELF, another inhibitory factor is the tripartite motif-containing 28 (known as TRIM28, KAP1, TIF1β), which has been shown to promote promoter-proximal pausing at cellular gene promoters [39, 58–60]. TRIM28 was initially identified as a transcriptional corepressor due to its interaction with members of the Kruppel transcription factor family (KRAB) and its potential direct binding to specific DNA sequences [58, 59]. These transcription factors often function as transcriptional repressors. When TRIM28 binds to KRAB-containing transcription factors, it facilitates the recruitment of co-repressors, histone deacetylases (HDACs), and chromatin remodeling complexes. This results in the compaction of chromatin structure and inhibition of gene transcription. In many inactivated genes, TRIM28 stabilizes the pausing of RNAP II near the transcriptional start site (TSS), which promotes promoter-proximal pausing and accumulation of RNAP II near gene promoter [58]. The modulation of RNAP II pausing depends on phosphorylation of TRIM28 at the specific site, Ser824. Similar to the SPT5 subunit of DSIF, the phosphorylation of TRIM28 is crucial in converting it from a pausing or negative elongation factor to a positive elongation factor [39, 58, 59, 61]. DNA-PK is the principal kinase that directly interacts with TRIM28 and catalyzes the phosphorylation of TRIM28 at serine 824 residue (p-TRIM28, S824), converting it to an elongation factor [39, 60]. However, pertaining to HIV transcription, the role of TRIM28 is still not clear. Nevertheless, TRIM28 is known to play a complex role in the control of HIV and other DNA/RNA viruses, influencing both positive and negative regulatory pathways. Specifically concerning HIV-1, TRIM28 is implicated in the regulation of viral latency and reactivation. However, further investigation is required to delineate its direct or indirect impact on HIV proviral gene expression. Initially, TRIM28 was identified as a restrictor of HIV through its interaction with Integrase, hindering viral integration into the host chromatin [62]. This discovery suggests that TRIM28 may functionally link integration and transcription processes. Subsequently, Randolph et al. [63] proposed a paradigm wherein TRIM28 governs a switch from repression to activation. Viruses could exploit a transcriptional repressor like TRIM28 for their activation by promoting site-specific phosphorylation (pS473 and/or pS824), thereby enhancing viral gene expression for infection and modulating immune gene expression for precise cell fate responses. Reports also suggested that TRIM28 also contribute to HIV-1 transcriptional inhibition by depleting Tat in myeloid lineage with the help of CTIP2 [64]. Consequently, targeting TRIM28 presents a promising therapeutic avenue during viral infection or latency by addressing upstream TRIM28 regulators, modulating TRIM28 enzymatic activities, and disrupting TRIM28 protein-protein interactions [63].

The elongation phase of HIV transcription is greatly enhanced by the Tat protein of HIV, as Tat enhances the recruitment of host cell elongation factor positive transcriptional elongation factor b (P-TEFb) to the HIV LTR. Subsequently, the CDK9 subunit of P-TEFb catalyze the hyper phosphorylation of RNAP II CTD at Ser2 and make RNAP II processive [49, 65]. In addition, CDK9 also catalyze the phosphorylation of negative factors, namely DSIF and NELF, and relieve their negative impact on HIV transcription [66, 67]. Our previous studies have shown that the lack of P- TEFb in quiescent primary T cells is responsible for HIV latency, even in the presence of adequate NF-kB activation [68]. P-TEFb complex consists of other subunits, mainly ELL2, ENL, AFF4, AF9; together, it is called super elongation complex (SEC) [69–71]. Thus, the HIV Tat protein plays a significant role in augmenting the elongation phase of HIV transcription and generating full-length genomic transcripts of HIV [1, 72–74]. In the absence of Tat, the elongation or completion of HIV transcripts is inefficient. Once HIV Tat is available, it positively regulates HIV transcription. Tat binding to trans-activation response (TAR) element, an RNA stem loop structure of HIV transcript, brings an essential transcriptional component, positive transcription elongation factor b (P-TEFb), thereby enhancing the efficiency of viral transcription. HIV transcription auto accelerates its transcription by generating more Tat protein [75, 76]. Thus, the enhanced rate of HIV transcriptional elongation results in a higher number of complete genomic HIV transcripts and generation of more viral particles.

In our previous publication, we clarified the important role of DNA-PK during HIV transcription and documented the continuous presence and gliding of DNA-PK with RNAP II along the HIV genome during transcription [37, 39]. Additionally, we identified the impact of cocaine use on promoting HIV transcription and replication [16, 17, 27, 28]. Later, we endeavored to define the underlying molecular mechanism through which cocaine augments HIV transcription and found that cocaine promoted HIV transcription by inducing different mechanisms [27, 28]. To expand upon this subject, in the present study, we focus on understanding the role of cocaine-stimulated DNA-PK in relieving RNAP II pausing during HIV transcription by catalyzing TRIM28 phosphorylation selectively at S824 residue. We found that cocaine further enhanced the nuclear localization of DNA-PK, where DNA-PK facilitates HIV transcription. We noted that cocaine exposure not only augmented the nuclear translocation but also enhanced its functional activity by increasing its phosphorylation at specific residue, Ser2056. Subsequently, we substantiated increased HIV transcription following cocaine exposure by examining the effect of cocaine-induced DNA-PK on the phosphorylation of specific sites on RNAP II CTD, namely Ser2 and Ser5. To further authenticate the precise role of cocaine-induced DNA-PK in CTD phosphorylation, we investigated the inhibitory potential of clinically evaluated DNA-PK inhibitors in reversing the influence of DNA-PK. These findings were further validated by conducting DNA-PK knockdown experiments in the presence or absence of cocaine, demonstrating the specific impact of cocaine-induced DNA-PK stimulation. Overall, our data demonstrate the crucial role of cocaine-mediated DNA-PK stimulation in relieving RNAP II pausing by converting TRIM28 from a transcriptional inhibitor to transcriptional activator protein. These findings are validated across diverse cell types belonging to both lymphoid and myeloid lineages, including microglia, the macrophages that reside in the CNS. This comprehensive study expands our understanding of the complex interplay among cocaine, DNA-PK, and TRIM28 and their influence on HIV transcription. Consequently, it illuminates potential therapeutic strategies for addressing HIV replication and/or mitigating the toxicities associated with drug abuse. Additionally, given that ART is unable to restrict HIV transcription or latency-reactivation, defining all factors and mechanisms that regulate HIV transcription will help open new avenues for better translational interventions.

## 2. Materials and Methods

### 2.1. Plasmid construction, gene transfer, transfection, and VSV-G pseudotyped virus generation

The pHR’p-Luc plasmid was constructed by inserting the EcoRI and XhoI fragment of HIV pNL4-3 into the pHR’ plasmid, as detailed previously [77, 78]. The procedure to knockdown the DNA-PK was also described previously [39]. The short-lived variant of green fluorescent protein (d2EGFP) was inserted at the nef position using the MluI and XhoI sites. Site-directed mutagenesis was conducted to substitute histidine at position 13 with leucine (H13L) (CAT to TTA), following established procedures [79, 80]. Human Embryonic Kidney 293 cells (HEK 293T) were cultured in Dulbecco’s Modified Eagle Medium (DMEM) supplemented with 2.05 ml-glutamine (Hyclone, ThermoScientific), 10% fetal bovine serum (Gemini), and 1 U/mL penicillin/streptomycin. Cells were seeded, grown to 70% confluency, and rinsed with Opti-MEM I (1X) + GlutaMAX-I Reduced Serum Medium (Gibco) before transfection. Transfection was done by using Lipofectamine 2000 (Invitrogen) as per the manufacturer’s instructions. Briefly, 35 µL of Lipofectamine 2000 reagent was mixed with 500 µL Opti-MEM. Separately, 18 µg of plasmid DNA mixture (3 µg pCMVΔ8.9.1, 4 µg pMD.G, 3 µg pMDL-g/p-RRE, 1 µg pRSV-Rev, and 7 µg of either pHR’P-Luc or pNL4-3-ΔE- EGFP for generating pNL4-3-ΔE-EGFP and pHR’p-P-Luc pseudotyped viruses, respectively) was prepared [28]. The two solutions were combined and incubated at room temperature (RT) for 30 minutes (min) to form the lipid-DNA complex, which was then introduced into the cells. Five hours after transfection, the culture medium was replaced with fresh DMEM. The cell supernatant containing the virus was collected at 48 hours (h) and 72 h post-transfection.

### 2.2. Generation of Luciferase cell line and latently infected Jurkat T-cell clones

The pHR’p-Luc virus was transduced into the Jurkat cell line via spinoculation in the presence of 8 µg/ml polybrene. Successful infection was subsequently confirmed by Luciferase assay [78]. The isolation of Clone 2D10 cells, characterized by the H13L Tat mutation, was detailed in our previous study [80]. Specifically, Vesicular Stomatitis Virus Protein G (VSV-G)-pseudotyped HIV particles were generated through triple transfection of 293T cells using Lipofectamine 2000 reagent (Invitrogen, Waltham, MA, USA). Virus titers were determined by infecting 2x10^6^ Jurkat T-cells with serial dilutions of concentrated virus preparation obtained from harvested medium supernatant. Six hours post-infection, cells were rinsed with phosphate-buffered saline (PBS), and RPMI 1640 medium was replenished. Expression of d2EGFP was assessed by fluorescently activated cell sorting analysis (FACS Calibur) 72 h post-infection, and d2EGFP expression was subsequently analyzed every week until cells were fully shut down without detectable d2EGFP expression before reactivation experiments.

### 2.3. Cell culture and cell experiments

Microglial, THP-1, MT-4, peripheral blood mononuclear cells (PBMC), Jurkat, and derivatives of Jurkat cells (Clone 2D10 and Jurkat-pHR’P-Luc) were cultured in either DMEM or RPMI 1640 medium. The culture medium was supplemented with 10% fetal bovine serum (FBS), penicillin (100 IU/ml), streptomycin (100 IU/ml), and 25 mM HEPES. Cells were maintained at 37°C in a 5% CO_2_ environment. Fresh medium was replenished every 2-3 days, and cell density was kept at 2x10^6^ cells/ml.

### 2.4. HIV replication-competent Virus

The Human Immunodeficiency Virus Type 1 (strain 93/TH/051) was obtained from the National Institute of Health AIDS reagent program. Primary HIV isolates were cultured following the instructions provided in the datasheet obtained through the UNAIDS Network for HIV Isolation and Characterization. Briefly, 4 X 10^6^ stimulated Jurkat cells (cells previously stimulated with PHA for 4 days and treated with polybrene) were collected and exposed to HIV (strain 93/TH/051) for 30 min at 37°C. Following this, fresh media was added, and the cells were incubated for 5 days. Cell free virus was recovered, aliquoted in multiple stock, and stored at -80°C till use.

### 2.5. Cocaine treatment and Inhibitor treatment

Cocaine hydrochloride was obtained from the National Institute on Drug Abuse (NIDA) Drug Supply Program. In this study, various cocaine concentrations were employed. Nonetheless, the maximum concentration utilized was 30 µM cocaine, which falls below the levels typically observed in the plasma of human drug users. All cocaine treatments were conducted at a concentration of 10 μM unless otherwise specified. Acute treatment involved exposing the cells to cocaine for 3 h, whereas chronic treatment entailed exposing the cells to cocaine twice daily for two consecutive days, with an additional 3h exposure prior to cell harvesting. Inhibitors (M3814 and NU7441) were treated for overnight (24 h) prior exposing to cocaine.

### 2.6. Infection of cells with replication-competent virus

Cells (5x10^6^ cells) were either untreated or exposed to cocaine for 3 h in the presence or absence of M3814 and, subsequently, were either uninfected or infected with replication-competent virus (1 mL) for 24 h and 48 h to assess HIV gene expression. Inhibitors were administered 24 h before HIV infection, with the specific doses mentioned in the figure legends.

### 2.7. Western blot analysis of total cell lysate

Cells (1x10^6^ or 5X10^6^ cells approx.) were treated with cocaine in the presence or absence of M3814 (DNA-PK inhibitor) and/or infected with a replication-competent virus for 24 h and 48 h. Subsequently, samples were collected and washed with 1 mL of ice-cold PBS, and 100 µL of 1X passive lysis buffer (Promega, Madison, WI, USA) was added to the cells. The cell lysate with the lysis buffer was then incubated on ice for 30 min. During the incubation, cells were vortexed for 30 seconds (sec) for complete lysis after every 10 min. Following incubation, the cell lysate was centrifuged at the highest speed for 30 min, and the supernatant was analyzed for protein concentration using the Pierce™ BCA Protein Assay Kit. Protein concentration was normalized, and an equal amount of protein was mixed with 5X Laemmle Sample buffer, heated to 95°C for 10 min, and then resolved by SDS-PAGE on a 6% or 12% gel at 120 volts until the dye reached the bottom. The resolved proteins were then transferred to a nitrocellulose membrane. The membranes were blocked with 3% Bovine serum albumin (BSA) for 1 h and incubated with primary antibodies at 4°C overnight and then with secondary antibody (1:15000 dilution) for 1 h at room temperature. After three washes with 1X TBST, the blot was detected using the Odyssey infrared imaging system application software version 3.0 (Li-cor Bioscience).

### 2.8. Western blot analysis of cytoplasmic and nuclear extracts

Cells (5x10^6^ or 1X 10^7^ cells approx.) were exposed to cocaine at various doses and time points, with or without the inhibitor. Subsequently, cells were collected and washed with 1 ml of ice-cold PBS. Following our established protocol, we fractionated cytosolic and nuclear proteins. Initially, cells were allowed to swell in 200 µl - 500 µl of cytoplasmic extract (CE) buffer (1 mM Hepes KOH pH 7.9, 60 mM KCl, 1 mM EDTA, 0.5% NP-40, 1 mM DTT, and 1 mM PMSF) for 10 min on ice, during which cells were vortexed for lysis. Nuclei were then pelleted at 4000 r.p.m for 10 min. The cytoplasmic lysates were transferred to new Eppendorf tubes for analysis of cytoplasmic proteins. The nuclei were washed twice with 1 ml of CE buffer, pelleted at high-speed centrifugation for 2 min, and subsequently resuspended in 80 µl of nuclear extract (NE) buffer (250 mM Tris pH 7.8, 60 mM HCl, 1 mM EDTA, 1 mM DTT, and 1 mM PMSF). The nuclei were lysed by 8 freeze-thaw cycles in liquid nitrogen. The nuclear lysate was cleared by centrifugation at high speed for 1 min, and the supernatant was transferred into a new microfuge tube. Total nuclear protein concentration in the samples was normalized using a standard BCA assay. An equal amount of total nuclear samples was loaded and resolved by 6% or 10% or 12% SDS-PAGE gel for electrophoresis. The proteins on the gels were transferred onto nitrocellulose membranes; blocked with 3% BSA for an hour; incubated with primary antibodies overnight and with secondary antibodies for an hour; and finally detected using the Odyssey infrared imaging system application software version 3.0 (Li-cor Bioscience).

### 2.9. Chromatin Immunoprecipitation (ChIP) assay

The ChIP assays were performed using our well-established protocol [81]. Briefly 1 x 10^8^ cells underwent fixation in 0.5% formaldehyde for 10 min with rotation at room temperature, facilitating the crosslinking of proteins to DNA. Subsequently, glycine was added to reverse the crosslinking process. Cells were harvested, washed twice with ice-cold PBS, and allowed to swell for 10 min in 5 ml CE Buffer. Nuclei were pelleted after centrifugation at 4000 rpm for 10 min and resuspended in 1 ml of SDS Lysis buffer (50 mM Tris-HCl, 1% SDS, 10 mM EDTA, 1 mM PMSF, 1 µg/ml aprotinin, 1 µg/ml pepstatin A). Genomic DNA was fragmented to lengths less than 800 bp by sonication (Misonex 3000) under the following conditions: Output 2.5 for 20 sec, repeated eight times. For each sample, 200 µl of sonicated samples were mixed with 800 µl of ChIP dilution buffer (0.01% SDS, 1.1% Triton X-100, 1.2 mM EDTA, 16.7 mM Tris-HCl pH 8.1, 167 mM NaCl). Samples were incubated with specific antibodies including IgG, DNA-PKcs, RNAP II, CDK7, CDK9, pTRIM28 (S824), and H3K27me3 at +4°C overnight. Protein A/G Sepharose was pre-saturated with salmon sperm DNA and 1% BSA, and 100 µl of 25% Protein A-Sepharose were utilized in DNA-protein immunoprecipitation. Following 3 h of incubation, Antibody-DNA-protein complexes were washed with 1 ml of each washing buffer. The first wash occurred with low salt immune complex wash buffer (0.1% SDS, 1% Triton X-100, 2 mM EDTA, 20 mM Tris-HCl pH 8.1, 150 mM NaCl), followed by high salt immune complex wash buffer (0.1% SDS, 1% Triton X-100, 2 mM EDTA, 20 mM Tris- HCl pH 8.1, 500 mM NaCl). The complexes underwent further washing with lithium chloride buffer (0.25 M LiCl, 1% NP-40, 1% sodium deoxycholate, 1 mM EDTA, and 10 mM Tris HCl pH 8.0) and twice with TE buffer (10 mM Tris-HCl pH 8.0, 1 mM EDTA pH 8.0). Protein DNA complexes were eluted from protein A/G Sepharose twice using 250 µl of freshly prepared elution buffer (1% SDS and 0.1 mM NaHCO3). Twenty microliters of 5 M NaCl were added to the total eluate, and Protein- DNA complexes were reversed-cross-linked at 65°C overnight. Ten microliters of 0.5 M EDTA, 10 µl of 2 M Tris-HCl pH 6.5, and 2 µl of 10 ng/ml proteinase-K were added, and samples were incubated at 50°C for 2 h followed by phenol extraction and ethanol precipitation. Precipitated DNA samples were dissolved in 100 µl of TE buffer, and 2 µl of the sample was utilized in real-time PCR using SYBR green PCR master mix (Thermo Scientific), following the method described previously by Kim et al [51]. No-antibody control values were subtracted from each sample value to eliminate non-specific background signal. The primer sets utilized in real-time PCR amplification are listed in Supplementary Table S1.

### 2.10. RNA extraction and real-time quantitative PCR (qPCR)

Total RNAs were extracted from 5×10^5^ cultured cells using an RNA isolation kit (Qiagen, Hilden, Germany) according to the manufacturer’s instructions. The isolated RNAs were meticulously assessed for their integrity, purity, and yield. Subsequently, using the isolated RNAs as the template, first-strand complementary DNA (cDNA) was synthesized utilizing M-MLV Reverse Transcriptase (Thermo Scientific, Waltham, MA). In brief, approximately 3 µg of extracted RNA was reverse transcribed in a total volume of 20 µl with 350 µM dNTP, 50 µM oligo (dT), 5X M-MuLV buffer, 200 U RNase inhibitors, and 200 U M-MuLV reverse transcriptase. The RNA, oligo (dT), and dNTPs were mixed and incubated at 65°C for 5 min, followed by 37°C for 50 min and 70°C for 10 min. The cDNA was subsequently diluted and subjected to real-time PCR using the Real-Time PCR system 7500TH (Life Technologies, Carlsbad, CA, USA). For all samples, Actin/GAPDH was measured as the internal control and utilized for data normalization. The primer sets utilized for the amplification are listed in Supplementary Table S1.

### 2.11. Luciferase assay

1x10^4^ or 5×10^5^ cells harboring pHR’P-Luc were plated in 12-well plates with complete RPMI media (supplemented with 10% FBS, penicillin, and streptavidin). The cells were incubated with cocaine (chronically, treating twice per day with cocaine) for 48 h in presence and absence of M3814. Luciferase levels in the cells were assessed using a Luciferase Assay System kit (Promega, Madison, WI, USA). Briefly, the cells were harvested, washed, and lysed with 1 X passive lysis buffer. After incubating 30 min at RT, cells were centrifuged at high speed for 2 min, and supernatant were transferred to a new Eppendorf tube. 10 µl of each sample lysate was added followed by 50 µl of luciferase substrate/assay buffer to individual wells of white plates to reflect light and maximize light output signal. Each sample was tested in triplicate. Luminescence was read in a Veritas Microplate Luminometer (Turner Biosystems).

### 2.12. Flow cytometry (FACS) analysis

FACS analyses were performed on 2D10 cells (Jurkat cells infected with VSV-G pseudotyped HIV virus carrying the GFP gene under the control of the HIV LTR promoter). Briefly, 2D10 cells were treated with inhibitor M3814 for 24 h. The next day, cells were activated/stimulated with 10 ng/ml Tumor Necrosis Factor alpha (TNF-α) for another 48 h. Cells were then harvested, washed, re- suspended with PBS, and analyzed with a FACS Calibur (BD Biosciences) using FlowJo software (Treestar Inc.).

### 2.13. Quantification and statistical analysis

Data are expressed as the mean standard deviation (mean ± SD). Comparisons between two groups were performed using Student’s t-test. Comparisons between more than two groups were carried out by one-way or two-way analysis of variance (ANOVA). If the p-value obtained from ANOVA was under 0.05 (p < 0.05), it was considered statistically significant. All statistical calculations were carried out using a GraphPad prism. All the statistical details of experiments can be found in the figure legends.

## 3. Results

### 3.1. Cocaine enhances both the catalytic activity and nuclear translocation of DNA-PK

The crucial role of DNA-PK during DNA double strand break repair is well established [43–45]. However, for the first time, we documented the vital role of DNA-PK in supporting gene transcription [37]. To define the underlying molecular mechanism through which DNA-PK augments HIV transcription, we confirmed that DNA-PK augment HIV transcription by supporting both the initiation and elongation phases of transcription [46]. Later, the crucial role of DNA-PK in supporting other cellular genes by enhancing RNAP II CTD phosphorylation was confirmed by us and others [39, 58–60]. Previously, we identified the significant impact of cocaine on enhancing HIV transcription and replication [27, 28]. However, to develop therapeutic strategies aimed at mitigating the toxic effects resulting from HIV replication and cocaine exposure, it is imperative to elucidate all the factors and mechanisms through which HIV and cocaine collaborate to induce cell toxicity via heightened HIV transcription. To investigate the role of cocaine in enhancing HIV transcription, we assessed the expression and nuclear level of DNA-PKcs, the catalytic subunit of DNA-PK. The impact of cocaine on the functional/catalytic activity of DNA-PK was evaluated by examining phosphorylation of p-DNA-PKcs at serine 2056 (p-DNA-PK S2056), a post translational modification that marks functionally active form of DNA-PK. We treated Jurkat cells, a T cell line, with increasing doses of cocaine for a duration of 3 h. Later, cells were harvested, and nuclear lysates were subjected to immunoblotting using antibodies specific for either total DNA-PKcs or phosphorylated form of DNA-PKcs (pDNA-PKcs S2056) to evaluate cocaine impact. We found higher levels of both DNA-PKcs and pDNA-PKcs S2056 in the nucleus upon cocaine exposure compared to the untreated cell control (Ctrl) (**Figures 1A & 1B**). The densitometric analyses of the protein bands validated a significant increase in the expression and nuclear level of both DNA- PKcs and pDNA-PKcs S2056 following cocaine-mediated cell stimulation. We further confirmed the effect of cocaine in upregulating and activating the DNA-PK in a dose-dependent manner using varying cell types belonging to different lineages, namely, microglial cells, a primary immune cell found in CNS and MT-4 **(Figures 1C, 1D, 1E & 1F).** These findings confirmed significant upregulation of DNA-PK expression and functional activation of DNA-PK (pDNA-PKcs S2056) by cocaine and in a cell lineage independent manner.

**Figure 1:**
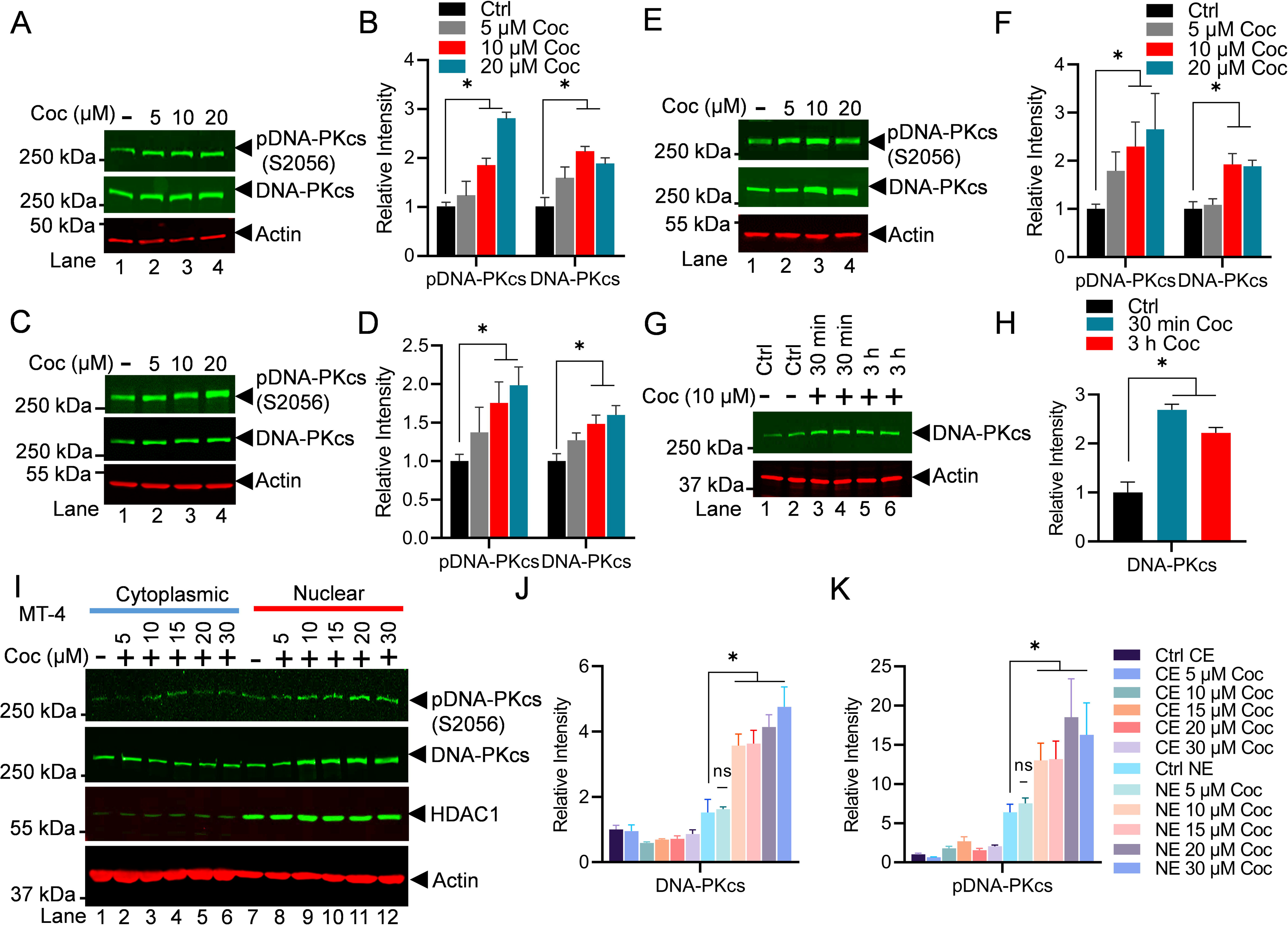
Cocaine enhances both the catalytic activity and nuclear translocation of DNA-PK. Jurkat cells harboring the pHR’-P-Luc provirus (**A**), microglial cells (**C**), and MT-4 cells (**E**) were treated with different concentrations of cocaine (Coc: 5, 10, and 20 μM) for 3 h (Lanes 2 to 4). Jurkat-pHR’-P’-Luc cells were reated with 10 µM cocaine (Coc) in replicates for 30 min and 3 h (Lanes 3 to 6) (**G**). Cells were harvested, and nuclear lysates were analyzed by immunoblotting using specific antibodies, pDNA-PKcs (S2056) and DNA-PKcs, as indicated. Actin, a constitutively expressed protein, was used as a loading control. Densitometric analysis of protein bands (normalized to actin) confirmed the significant upregulation of total DNA-PKcs and its phosphorylated form, pDNA-PKcs S2056 (pDNA-PKcs), following cocaine treatment (**B, D, F, & H**). MT-4 cells were treated with increasing doses of cocaine for 3 h. Cells were harvested and lysed, and both cellular and nuclear lysates were analyzed by immunoblotting with antibodies against pDNA-PKcs (S2056), DNA-PKcs, HDAC1, and Actin (**I**). Densitometric analysis of protein bands, normalized to actin, validated the enhancement in both the catalytic activity and nuclear translocation of DNA-PK (**J & K**). mmunoblots are representative of at least three independent experiments. The results are expressed as mean ± SD and analyzed by one- or two-way ANOVA, followed by Tukey’s multiple comparison test. Asterisks over the bars indicate significant differences: ∗p < 0.05 for the comparison of cocaine-treated cells vs. untreated cells (Ctrl).

Subsequently, we examined the impact of cocaine on DNA-PK levels and activation in a time- dependent manner (**Figures 1G & 1H**) by treating the Jurkat cells infected with pHR’P-Luc with a fixed dose of cocaine (10 μM) for 30 min and 3 h, with untreated cells as a control (**Figures 1G & 1H**). Upon analyzing the nuclear extract, we found upregulation of nuclear DNA-PK level within 30 min, which remained higher even after 3 h.

Furthermore, to establish the ubiquity of the phenomenon, MT-4 cells were treated with increasing doses of cocaine for 3 h, and the translocation of DNA-PKcs from cytoplasm to nucleus was evaluated by immunoblotting, analyzing both cytoplasmic and nuclear protein fractions on the same blot. As a control, we evaluated HDAC-1 levels, a protein that predominantly exists in the nucleus, and only a small portion was present in the cell cytoplasm. Accordingly, we found abundant presence of HDAC-1 in the nuclear extract of the cell, validating the purity of our nuclear fraction and our assay conditions. As loading control, we examined the presence of Beta-actin protein, which is constitutively expressed in the cell and can be detected in both cytoplasmic and nuclear fractions. Interestingly, we noted significantly enhanced translocation of both DNA-PKcs and (pDNA-PKcs 2056) into the nucleus following cocaine treatments (**Figures 1I, 1J & 1K**). The enhanced nuclear localization of DNA-PK following cell stimulation was also observed previously [38]. These results suggest that cocaine augments DNA-PK function both by enhancing its upregulation and nuclear translocation, besides augmenting the catalytic activity of DNA-PK by specifically increasing its phosphorylation at S2056. Moreover, higher nuclear translocation of DNA-PK following cocaine exposure clearly suggests a role for DNA-PK in DNA transections, including transcription. Altogether, these results confirm that cocaine intake promotes activation of DNA-PK by enhancing both the nuclear translocation and functional activity of DNA-PK.

### 3.2. Cocaine-induced HIV transcription augments overall HIV replication

To evaluate the impact of cocaine on HIV transcription and subsequently to HIV gene expression, we freshly infected Jurkat cells with non-replicating attenuated HIV, pHR’P-Luc, to generate the Jurkat-pHR’P-Luc cell line [78]. The pHR’P-Luc is an HIV-based lentivirus that expresses luciferase reporter gene under the control of the HIV LTR promoter (**Figure 2A**). Therefore, expression of luciferase indicates ongoing HIV transcription and gene expression. **Figures 2B and 2D** depict the schematic overview of the cell treatment procedures. As anticipated from our previous studies [27, 28], a significant increase in luciferase counts was observed in a dose-dependent manner, validating cocaine-mediated upregulation of HIV gene expression (**Figure 2C**). To further confirm the impact of cocaine-mediated cell stimulation on HIV gene expression and replication, PBMC cells were chronically treated with cocaine prior to being infected with a replication-competent dual tropic HIV Type 1 strain 93/TH/051 for a period of 24 h. The HIV transcripts were quantified via real- time qPCR using primer sets that amplify the Envelope (*Env)* region of the HIV genome. A significant upregulation of HIV gene expression was confirmed in the presence of cocaine (**Figure 2E**). Next, The HIV protein expression was evaluated via immunoblotting using antibodies against Gag subunits (p24) of HIV by comparing the cell lysates of cocaine treated or untreated HIV infected cells (**Figures 2F & 2G**). The upregulation of p24 confirms enhanced HIV gene expression and replication in the presence of cocaine. Together, these results suggest that cocaine induced signaling pathways promote activation of both cell status and transcription machinery, including DNA-PK stimulation (p-DNA-PK S2056) (**Figure 1**), resulting in enhanced HIV transcription and consequently higher HIV replication.

**Figure 2:**
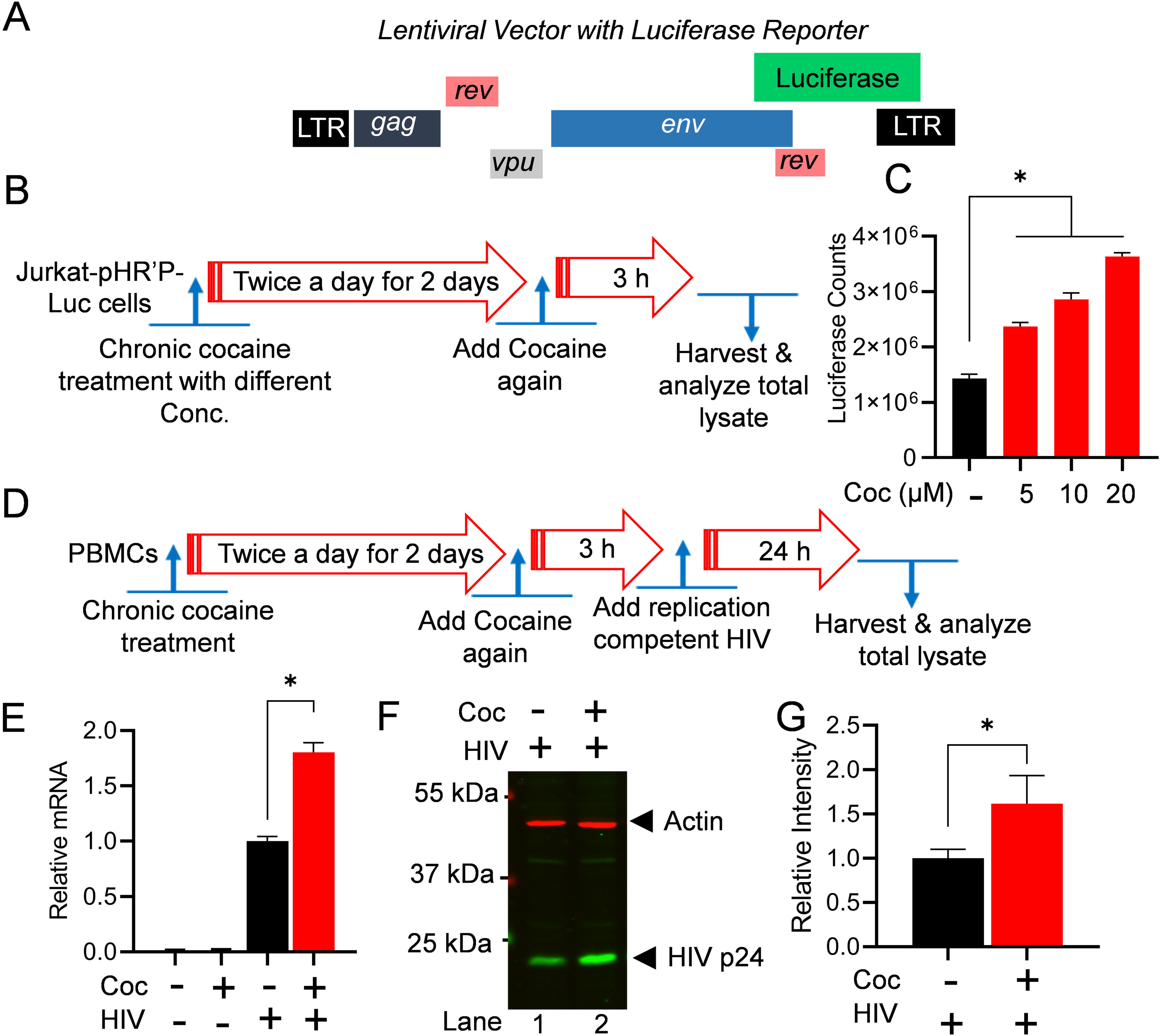
Cocaine-induced HIV transcription augments overall HIV replication. Structure of the lentiviral vector (pHR’-PNL-Luc) carrying the reporter luciferase gene under the HIV LTR promoter (**A**). Schematic representation of the cocaine (Coc) treatment for the luciferase reporter assay (**B**). Jurkat-pHR’-P-Luc cells were chronically treated with 5 µM – 20 µM of cocaine. The cells were lysed, and luciferase reporter protein expression levels were assessed using luciferase assays (**C**). Schematic depiction of the cocaine treatment and subsequent infection of PBMC cells with replication-competent HIV (**D**). HIV transcripts were quantified by real-time PCR using primer sets that amplify the Envelope (Env) region of the HIV genome (**E**). The level of Gag/p24 protein was analyzed by immunoblotting with specific antibodies against HIV p24 (**F**). Actin, a constitutively expressed protein, was used as a loading control in the same blot. Densitometric analysis of protein bands (normalized to actin) confirmed a significant increase in p24 levels compared to untreated cells (Ctrl) (**G**). Immunoblots are representative of at least three independent experiments. The results are expressed as mean ± SD, analyzed by one-way ANOVA followed by Tukey’s multiple comparison test (**C & E**) or unpaired t-test (**G**). Asterisks over the bars indicate significant differences: ∗p < 0.05 for the comparison of cocaine-treated cells vs. untreated cells.

### 3.3. Partial DNA-PK inhibition is sufficient to restrict HIV transcription, replication, and latency reactivation

We have shown that DNA-PK plays an important role during HIV transcription [37, 39]. To extend further on those findings and establish the translational potential of DNA-PK inhibition in restricting HIV transcription and replication, we evaluated the role of a clinically evaluated DNA-PK inhibitor (DNA-PKi), M3814. Interestingly, in a recent clinical study, DNA-PK inhibitors, including M3814 at dosages from 110 µM to 320 µM were found safe and highly effective as potential anti-cancer drugs [82–92], validating the safety of these agents for human use [84]. Notably, we found that partial DNA-PK inhibition by only 20 µM (less than 1/5) is sufficient to restrict HIV transcription, replication, and latency reactivation without any cell toxicity.

We assessed the effect of M3814 on HIV transcription and latency reactivation. The infected Jurkat cells that harbor latent HIV provirus (pHR’P-Luc) in their genome, which expresses luciferase reporter gene under the control of HIV LTR promoter (**Figure 2A**). These cells, Jurkat-pHR’P-Luc, were incubated overnight (24 h) with increasing concentrations (5 µM, 10 µM, 15 µM and 20 µM) of M3814. The next day, the cells were stimulated with 10 ng/ml Tumor Necrosis Factor alpha (TNF-α) for another 48 h (**Figure 3A**). A strong M3814-mediated dose-dependent inhibition of HIV transcription was observed, indicated by highly reduced luciferase counts, marking restricted HIV gene expression when DNA-PK was selectively inhibited using highly specific and clinically evaluated DNA-PKi (**Figure 3B**). As controls, cells were either treated with TNF-α alone (positive control) or left untreated (negative control). The inverse correlation between luciferase counts and M3814 concentration confirms a direct role DNA-PK in supporting HIV transcription and latency reactivation (**Figure 3B**). These findings were further validated by examining the presence of luciferase protein in the cell extracts by performing immunoblotting using antibody specific to Luciferase protein (Luciferase antibody: sc-74548) (**Figure 3C**). The strong dose-dependent inhibition of luciferase by M3814 established a vital role of DNA-PK during HIV transcription. Overall, these findings demonstrate a pivotal role of DNA-PK in supporting HIV transcription and latency reactivation. Moreover, the data obtained confirm our previous findings where we used another highly specific clinically tested DNA-PKi (Nu7441) [39].

**Figure 3:**
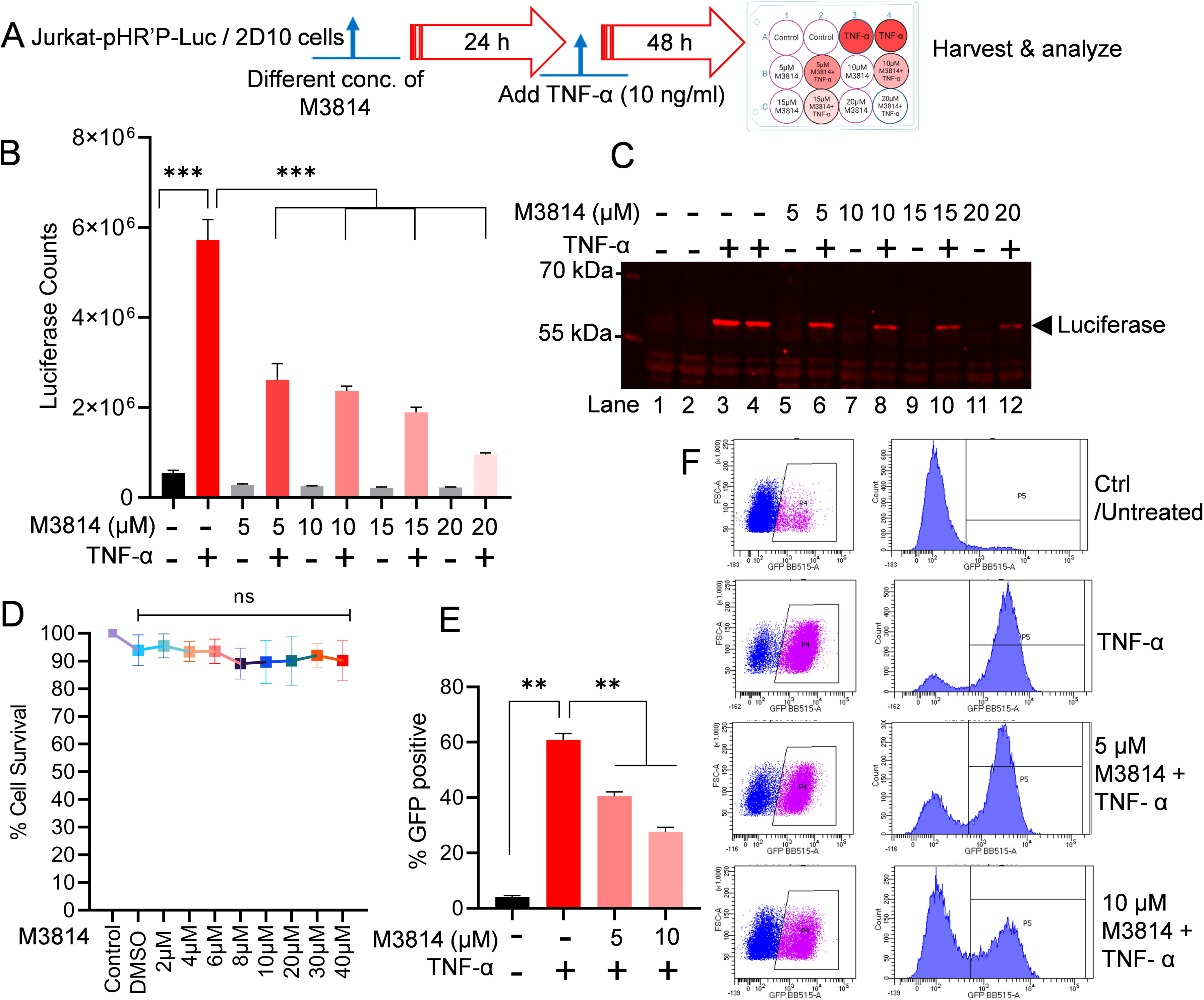
Partial DNA-PK inhibition severely impairs HIV transcription and latency reactivation. Schematic representation of protocol for M3814 inhibitor and TNF-α treatment in the luciferase reporter assay (**A**). Jurkat-pHR’-P-Luc cells were treated with 5, 10, 15, and 20 µM of M3814 for 24 h, followed by activation with TNF-α (10 ng/ml) for another 48 h. Cells were lysed, and the level of reporter protein expression was determined by a luciferase assay (**B**). The same lysates were analyzed by immunoblotting using specific antibodies against the luciferase protein (sc-74548) (**C**). Jurkat-pHR’-P-Luc cells were cultured with different concentrations (2 μM to 40 μM) of M3814 for 48–72 h, and cell cytotoxicity was determined via MTS-PMS cell proliferation assay (Promega, Madison, WI, USA) (**D**). Latently HIV-infected 2D10 cells, which express the reporter short-lived green fluorescent protein (d2EGFP) from the HIV LTR promoter, were treated with 5 µM or 10 µM of M3814 for 24 h and then stimulated with TNF-α for another 48 h. Cells were subjected to GFP expression analysis via flow cytometry (**E & F**). Immunoblots are representative of at least three independent experiments The results are expressed as mean ± SD and analyzed by one- or two-way ANOVA followed by Tukey’s multiple comparison test. Asterisks over the bars indicate significant differences: **p < 0.01 and ***p < 0.001 for the comparison of inactive vs. activated cells (TNF-α) and activated cells (TNF-α) vs. activated cells (TNF-α) in the presence of the DNA-PK inhibitor, M3814.

To exclude the possibility that the reduced luciferase activity upon M3814 treatment was not due to cell loss, we performed cell viability assay. The Jurkat-pHR’P-Luc cells were cultured with different concentrations (2 μM-40 μM) for M3814 for 48–72 h, and cell cytotoxicity was determined by MTS-PMS cell proliferation assay (Promega, Madison, WI, USA). We did not observe any significant cell cytotoxicity even at 40 μM of M3814 treatment (**Figure 3D**).

The impact of M3814 in restricting the reactivation of latent HIV was further confirmed using another latently infected cell line, 2D10 cells. The 2D10-cell line is a latently infected Jurkat T-cell line, which harbors a latent HIV provirus in their genome that expresses a reporter short-lived green fluorescent protein (d2EGFP) from HIV LTR promoter [78, 80]. Thus, GFP expression marks ongoing HIV gene expression. The 2D10 cells were treated for 24 h with different doses of M3814. Next day, cells were activated with 10 ng/ml TNF-α for another 48 h. Later, we quantified GFP expression through flow cytometric analysis. The TNF-α, which we used as a positive control, was able to stimulate latent HIV in more than 90% of cells compared to the control (unstimulated cells), marked by GFP expression in most (90%) cells. As anticipated, we observed a clear dose-dependent inhibition of HIV proviral reactivation upon DNA-PK inhibition, indicated by the reduced GFP expression in cells treated with the M3814 in a dose-dependent manner compared to the positive control (TNF-α treated) (**Figure 3E & 3F**). Overall, these data suggested that DNA-PK-mediated stimulation of HIV transcription is required for the reactivation of latent HIV provirus.

To assess the impact of different highly specific and clinically evaluated DNA-PK inhibitors on HIV replication, Jurkat cells were treated with the increasing doses of different DNA-PK inhibitors, M3814, and NU7441 for 24 h. The next day, cells were activated with 10 ng/ml TNF-α for 3 h. Later, cells were infected with a replication-competent dual tropic HIV (Type 1 strain 93/TH/051). The cell lysates were prepared after either 4 h post infection (4hpi) or 6 h post infection (6hpi), as shown in the figure (**Figure 4A**). The lysates were analyzed by immunoblotting with HIV cocktail antibodies p55, p24, and p17. The results show a clear inhibition of all HIV protein (HIV p55, HIV p24, and HIV p17) with increasing doses of DNA-PK inhibitors M3814 (**Figures 4B & 4C**) and NU7441 (**Figures 4D & 4E**). The stronger suppression of HIV replication was noted with the increasing doses of DNA-PK inhibitors, indicating the target-specific inhibition and confirming the vital role of DNA-PK-induced HIV transcription in supporting overall HIV replication. Additionally, the data confirm that in the presence of DNA-PKi, TNF-α mediated strong cell stimulation and NF-kB activation is ineffective in inducing HIV transcription, which suggests that not only cocaine but also TNF-α/NF- kB-mediated HIV transcription requires functional DNA-PK.

**Figure 4:**
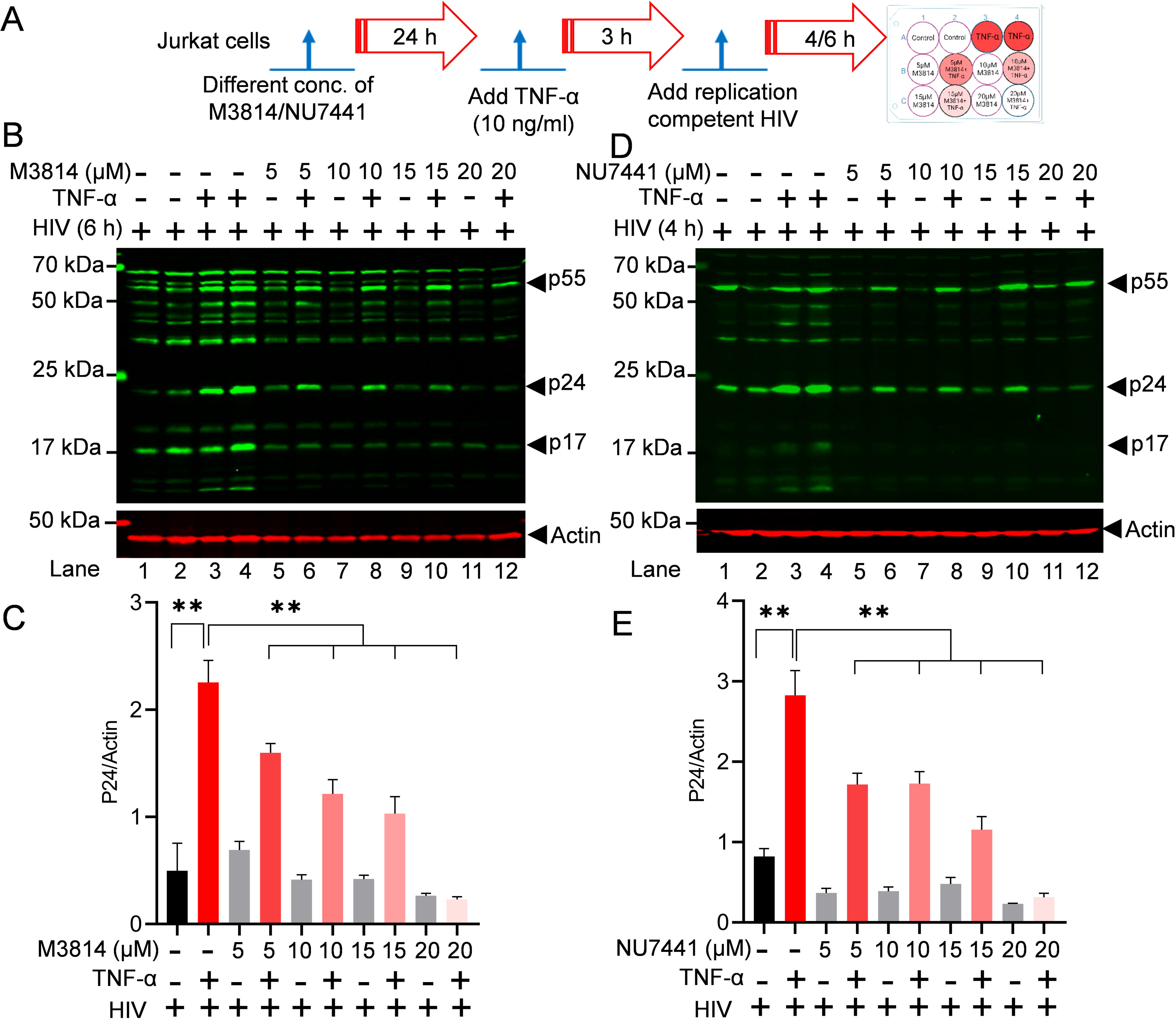
Partial DNA-PK inhibition restricts HIV replication. Schematic timeline for the treatment with M3814, NU7441 inhibitors, TNF-α, and replication-competent HIV (**A**). Jurkat cells were treated overnight with different concentrations of M3814 (5, 10, 15, and 20 μM) (**B**) and NU7441 (5, 10, 15, and 20 µM) (**D**) for 24 h (Lanes 5-12). The next day, cells were activated with 10 ng/ml TNF-α for 3 h (Lanes 3, 4, 6, 8, 10, & 12). Subsequently, cells were infected with a replication-competent dual-tropic HIV (Type 1 strain 93/TH/051) (Lanes 1-12). Cell lysates were prepared 4 h (NU7441) or 6 h (M3814) post-infection (hpi). Total cell lysates were analyzed by SDS-PAGE, transferred to a nitrocellulose membrane, and detected with specific HIV antibodies as indicated. Immunoreactive proteins were detected using appropriately labeled secondary antibodies with Licor. Actin was used as a loading control. Densitometric analysis of protein bands relative to actin (**C & E**). Immunoblots are representative of at least three independent experiments. The results are expressed as mean ± SD and analyzed by one-way ANOVA followed by Tukey’s multiple comparison test. Asterisks over the bars indicate significant differences: **p < 0.01 for the comparison of inactive vs. activated cells (TNF-α) and activated cells (TNF-α) vs. activated cells (TNF-α) in the presence of DNA-PK inhibitors, NU7441 or M3814.

### 3.4. DNA-PK inhibition strongly suppresses cocaine induced HIV transcription in primary cells, as well

The above data and our previous publication suggested that cocaine plays a significant role in enhancing HIV transcription and replication [27, 28]. In order to understand the molecular mechanisms by which cocaine controls HIV transcription and gene regulation, we investigated whether cocaine promotes HIV transcription and replication by enhancing both the catalytic activity and nuclear translocation of DNA-PK. To test this hypothesis, we treated the cells infected with pHR’P-Luc, which carry proviral HIV and expresses luciferase reporter under HIV LTR promoter, with 10 µM M3814 for 24 h. The next day, cells were treated with cocaine chronically for two days (10 µM cocaine twice a day for 3 days). Later, the cell extracts were prepared, and the level of luciferase reporter protein expression was determined via luciferase assays. As anticipated from the above analysis (**Figure 2B**), we noticed significant upregulation of luciferase counts, indicating enhanced HIV transcription in cocaine-treated samples. However, in the presence of M3814, HIV transcription is strongly restricted both in the presence and absence of cocaine (**Figure 5A**). These results confirmed the specific role of cocaine-stimulated DNA-PK in promoting HIV transcription and gene expression. Subsequently, to assess the effect of cocaine-mediated DNA-PK stimulation on HIV transcription and replication, we treated the Jurkat T cells (**Figure 5B & 5C**) and PBMC (**Figure 5D, 5E & Supplementary Figure S1**) with M3814 overnight (24 h). The next day, fresh media was provided with cocaine for 3 h. After 3 h of cocaine exposure, cells were infected with replication competent virus (93/TH/051) for 24 h. The HIV transcripts were quantified using real-time qPCR using primer sets that amplify the Nuc-2 (**Figure 5B & 5D**) and *Env* (**Figure 5C & 5E**) region of the HIV genome. A significant upregulation of HIV transcript was observed in the presence of cocaine, but, as anticipated, the presence of M3814 strongly restricted HIV gene expression in a dose-dependent manner (**Figure 5B, 5C, 5D & 5E**). These results were further validated by examining the expression of HIV protein in the absence or presence of M3814. The Jurkat cells were treated with M3814 for 24 h. The next day, cells were treated with cocaine for 3 h. Later, we infected the cells with replication competent HIV (93/TH/051) for another 24 h. The cell lysates were then subjected to immunoblotting to detect HIV proteins p24 and p17. Again, we noted higher levels of HIV proteins, p24 and p17, following cocaine exposure. However, in the presence of inhibitor, the level of HIV proteins dropped sharply, further demonstrating that cocaine-induced DNA-PK plays a crucial role in HIV transcription and replication (**Figure 5F & 5G**). Overall, these results confirm that cocaine-mediated DNA-PK stimulation is required for HIV gene expression and consequently for HIV replication.

**Figure 5:**
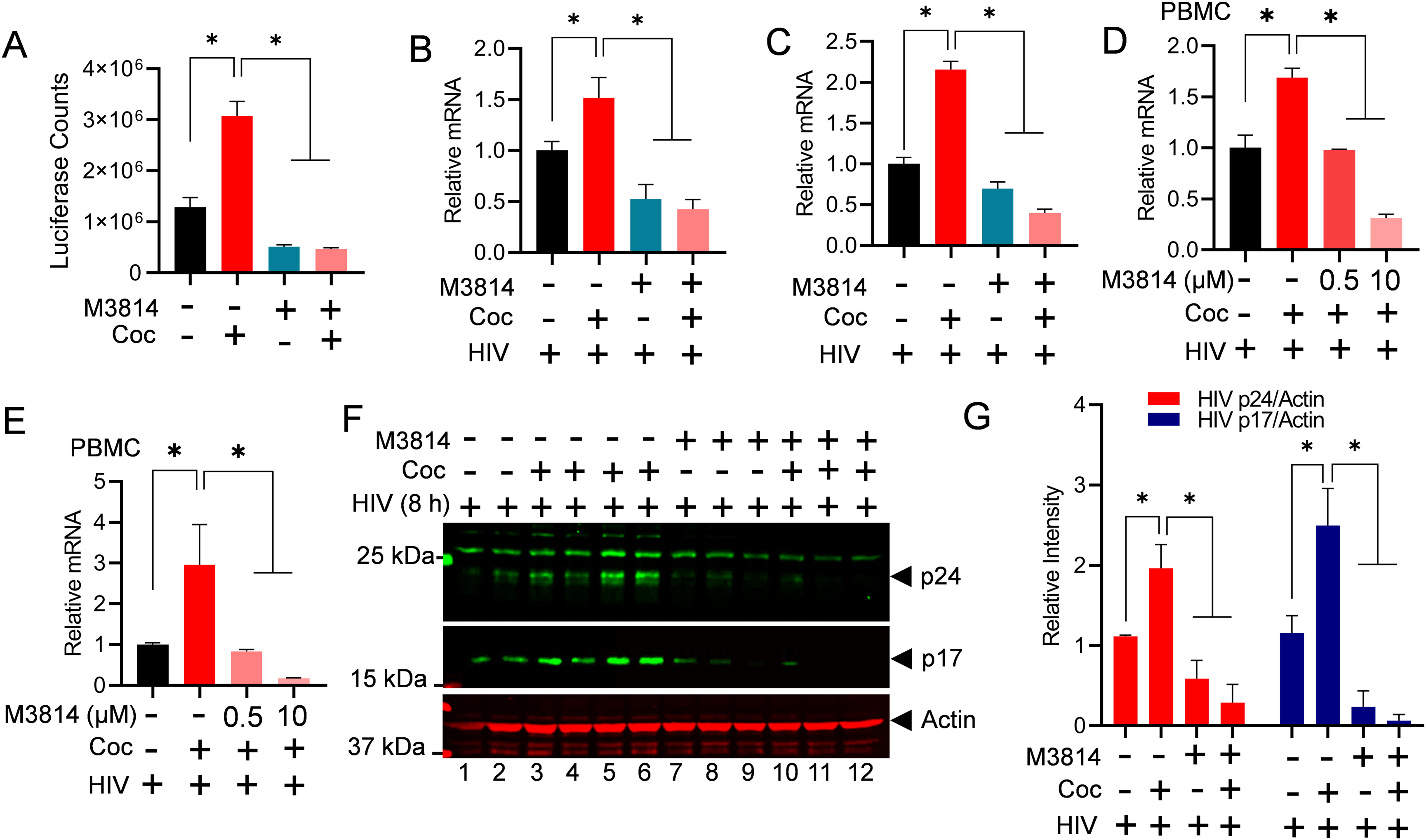
Cocaine-mediated DNA-PK activation promotes HIV transcription and replication in both cell ines and primary cells. Jurkat-pHR’-P-Luc cells were treated with 10 µM of M3814 for 24 h. The next day, cells were treated with cocaine twice daily for 48 h and again 3 h before harvesting. Cells were lysed, and the evel of reporter protein expression was determined using a luciferase reporter assay (**A**). Jurkat cells (**B & C**) and PBMCs (**D & E**) were treated with 10 µM of M3814 for 24 h, then treated with cocaine for 3 h, and subsequently infected with replication-competent HIV for another 3 to 6 h. HIV transcripts were quantified by real-time PCR using primer sets that amplify the Nuc-2 (**B & D**) and Env (**C & E**) regions of the HIV genome. Jurkat cells were treated with 10 µM of M3814 for 24 h (Lanes 7 to 12), then treated with cocaine for 3 h (Lanes 3-6 & 10-12) and infected with replication-competent HIV across all lanes (Lanes 1-12) for another 5 h. The levels of HIV p24 and p17 proteins were analyzed via immunoblotting using antibodies against these HIV proteins (**F**). Actin, a constitutively expressed protein, was used as a loading control. Densitometric analysis of protein bands (normalized to actin) was performed (**G**). Immunoblots are representative of at least hree independent experiments. The results are expressed as mean ± SD and analyzed by two-way ANOVA ollowed by Tukey’s multiple comparisons test. Asterisks over the bars indicate significant differences: ∗p < 0.05 for the comparison of cocaine-treated samples vs. untreated (Ctrl) and the comparison of cocaine plus nhibitor-treated samples vs. cocaine alone-treated samples.

### 3.5. Cocaine promotes HIV transcription by enhancing the phosphorylation of the C-terminal domain (CTD) of RNA polymerase II (RNAP II)

RNAP II is the main enzyme that transcribes eukaryotic DNA into mRNA. The C-terminal domain (CTD) of RNAP II consists of a repeating sequence of 7 amino acids (heptapeptide) with the consensus sequence Tyr1-Ser2-Pro3-Thr4-Ser5-Pro6-Ser7 (YSPTSPS) around 52 times [32–35]. All residues within the CTD heptad repeat can be post-translationally modified by phosphorylation (tyrosine, threonine, serine, and proline). However, in RNAP II CTD, Serine 5 and Serine 2 phosphorylation (Ser5-P and Ser2-P) are the best studied and the most established indicators of ongoing transcription. Specifically, the phosphorylation of RNAP II CTD at Ser5 is linked to the initiation phase of transcription, marking initial movement of RNAP II from the gene promoter, whereas phosphorylation of Ser2 is found to be correlated with the elongation phase of transcription. Notably, to generate a full-length HIV transcript, both initiation and elongation phases are required. Therefore, we evaluated if cocaine enhances HIV transcription by hyper-phosphorylating RNAP II CTD, we analyzed phosphorylation of RNAP II CTD at Ser2 and Ser5 upon cocaine exposure. THP-1 cells were treated with increasing concentrations of cocaine for 3 h. Later, nuclear lysate was subjected to immunoblotting to probe with RNAP ll Ser2-P, RNAP II Ser5- P, and RNAP II Total. The activation of p65, a subunit of NF-kB, was analyzed as a positive control to confirm cocaine-mediated cell stimulation. As anticipated, we observed stimulation of p65, marked by enhanced level of p65 in the nucleus compared to untreated cells (Ctrl). Notably, we also found hyper-phosphorylates RNAP II CTD at both Ser2 and Ser5 residues following cocaine treatment (**Figures 6A & 6B**). The dose-dependent upregulation of RNAP II CTD phosphorylation further confirmed the direct impact of cocaine in enhancing the phosphorylation of RNAP II CTD.

**Figure 6:**
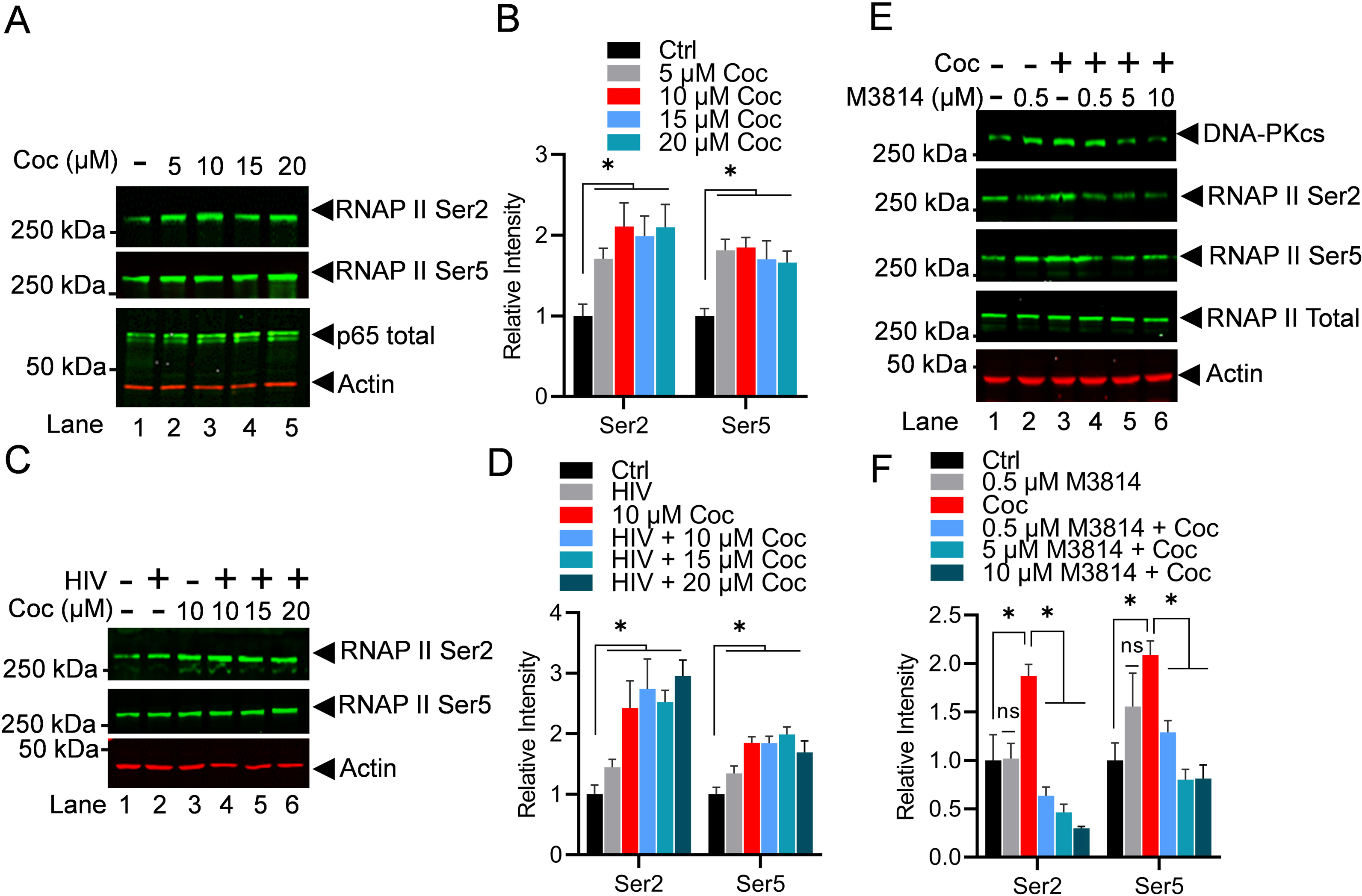
Cocaine promotes HIV transcription by enhancing the phosphorylation of the C-terminal domain (CTD) of RNA polymerase II (RNAP II). THP-1 cells were treated with increasing doses of cocaine (5, 10, 15, and 20 µM) for 3 h (**A**). MT-4 cells were treated as follows: untreated and uninfected (Lane 1), nfected with HIV (93/TH/051) without cocaine treatment (Lane 2), treated with cocaine without HIV infection (Lane 3), or pre-treated with different concentrations of cocaine before HIV infection (Lanes 4 to 6) (**C**). Cells were harvested, and nuclear lysates were analyzed by immunoblotting with specific antibodies against phosphorylated RNAP II, RNAP II Ser2, and RNAP II Ser5. Actin, a constitutively expressed protein, was used as a loading control. Densitometric analysis of protein bands (normalized to actin) confirmed significant hyper-phosphorylation of RNAP II CTD at both Ser2 and Ser5 residues following cocaine treatment (**B & D**). THP-1 cells were treated with cocaine in the absence or presence of different concentration of M3814 (0.5, 5, and 10 µM) (**E**). Cells were harvested, and nuclear extracts were evaluated via immunoblotting using specific antibodies against RNAP II Ser2, RNAP II Ser5, and total RNAP II. Densitometric analysis of protein bands (normalized to actin) confirmed a significant increase in RNAP II CTD phosphorylation at both Ser2 and Ser5 upon cocaine treatment. However, a significant reduction in CTD phosphorylation at both Ser2 and Ser5 was observed upon DNA-PK inhibition with M3814 compared to cocaine-alone samples (**F**). Immunoblots are representative of at least three independent experiments. The results are expressed as mean ± SD and analyzed by two-way ANOVA followed by Tukey’s multiple comparison test. Asterisks over the bars indicate significant differences. ∗ p < 0.05 is for the comparison of cocaine-treated samples against untreated (Ctrl) and the comparison of cocaine plus inhibitors treated against cocaine alone-treated samples

To further validate the ubiquity of our findings, results were confirmed in MT-4 cells. The cells were treated with different doses of cocaine for 3 h before being infected with a dual tropic HIV (93/TH/051). After 3 h, nuclear extracts were examined for RNAP II at Ser2 and Ser5. The hyper- phosphorylation of RNAP II at both the Ser2 and Ser5 positions of RNAP II CTD upon cocaine treatment was evaluated (**Figures 6C & 6D**). The dose-dependent hyper-phosphorylation of RNAP II CTD was clearly evident.

Subsequently, we examined if DNA-PK is involved in the cocaine-induced RNAP II CTD phosphorylation. We hypothesized that if cocaine-induced DNA-PK catalyzes the RNAP II CTD phosphorylation, then inhibition of DNA-PK should impair the cocaine stimulated RNAP II CTD hyper-phosphorylation. To test this hypothesis, the THP-1 cells were treated with increasing concentrations of M3814 for 24 h. Next day, cells were exposed to cocaine for 3 h. Later, nuclear protein lysates were analyzed by immunoblotting to examine RNAP II CTD phosphorylation at the sites Ser2 and Ser5. As shown in the figure (**Figure 6E**), cocaine treatment significantly upregulates RNAP II CTD phosphorylation at Ser2 and Ser5, validating the above results. We noted a significant reduction of CTD phosphorylation at both Ser2 and Ser5 in the presence of M3814 when compared to cocaine alone samples. The dose-dependent inhibition of RNAP II CTD phosphorylation at both Ser5 and Ser2 with M3814 confirmed our hypothesis and validated that cocaine-stimulated DNA-PK plays a vital role in promoting both the initiation and elongation phases of HIV transcription by catalyzing both Ser5 and Ser 2, respectively (**Figures 6E & 6F**). Overall, the results demonstrate that by activating DNA-PK, cocaine promotes different stages of HIV transcription, a necessity to produce complete HIV genomic transcripts or new HIV progeny.

### 3.6. Cocaine enhances the elongation phase of HIV transcription not only by stimulating DNA-PK but also via P-TEFb activation

The above results demonstrate that cocaine promotes both the initiation and elongation phases of HIV transcription by enhancing RNAP II CTD phosphorylation at Ser5 and Ser2. We further investigated if cocaine promotes the elongation phase by stimulating P-TEFb. The CDK9 is the kinase subunit of P-TEFb complex, which plays a crucial role in catalyzing the phosphorylation of RNAP II CTD at position Ser2, a post-translational modification that makes RNAP II processive or elongation proficient. We examined the stimulation of P-TEFb following cocaine exposure. Jurkat- pHR’P-Luc cells were exposed to increasing doses of cocaine for 3 h. Subsequently, nuclear lysates were subjected to immunoblotting using specific antibodies against CDK7 (TFIIH), p-CDK9 (thr186), and total CDK9. The data indicated that cocaine enhances CDK7, thereby facilitating the initiation of HIV transcription. Additionally, the data shows an increase in the phosphorylation of CDK9 at threonine residue 186, which marks functionally active CDK9. However, cocaine did not affect the level of total CDK9 (**Figures 7A & 7B**). To further validate these findings, Jurkat-pHR’P- Luc cells were treated with escalating doses of cocaine for 2 h and subsequently infected with replication-competent HIV (strain 93/TH/051) for an additional hour. As shown in the **Figure 7C**, Jurkat-pHR’P-Luc cells were treated as follows: untreated and uninfected (Lane 1), infected with HIV (93/TH/051) in the absence of cocaine (Lane 2), treated with cocaine without HIV infection (Lane 3), or pre-treatment with different concentrations of cocaine before infecting with HIV (Lane 4–6). The nuclear lysates were analyzed via immunoblotting using specific antibodies against main P-TEFb subunits, CDK9 and Cyclin T1. The obtained data clearly shows the enhanced phosphorylation of CDK9 and also upregulation of Cyclin T1 upon cocaine treatment, demonstrating that cocaine further supports the ongoing elongation phase of HIV transcription by stimulating CDK9. Nevertheless, it does not affect the level of CDK9. Actin was used as a loading control, while P24 was probed to mark the ongoing HIV replication. Densitometric analysis of protein bands validated a significant increase to p-CDK9 (thr186) and Cyclin T1 but not CDK9 total levels compared to untreated cells (control) (**Figures 7C & D)**. We also evaluated the impact of cocaine on another kinase, CDK7, which is a component of TFIIH complex that is mainly responsible for Ser5 phosphorylation, another RNAP II CTD modification required for the initiation phase of transcription. As expected, we noted upregulation of CDK7 upon cocaine treatment. The results again demonstrate that by enhancing CDK7, cocaine facilitates the initiation phase of HIV transcription.

**Figure 7:**
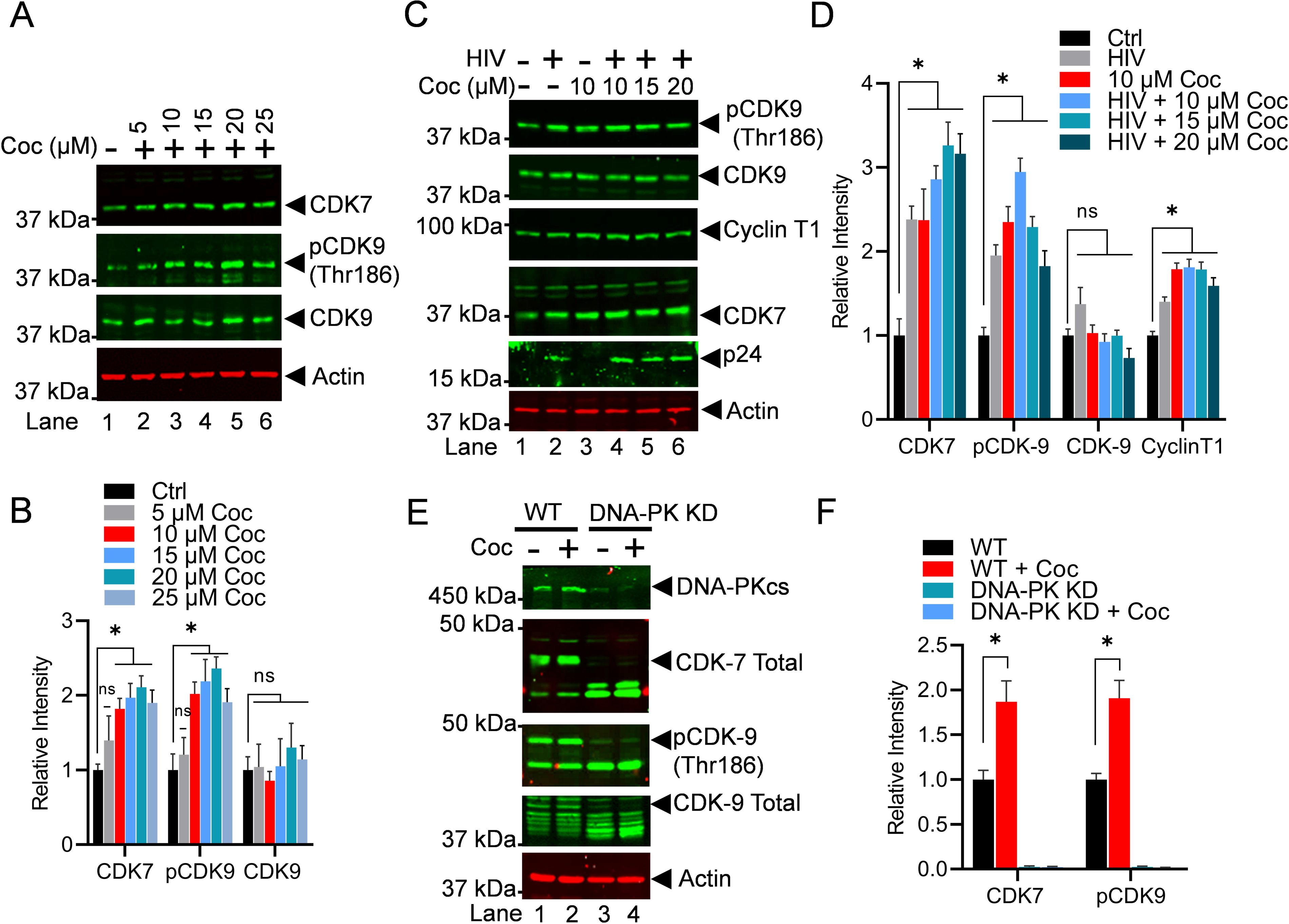
Cocaine enhances the elongation phase of HIV transcription not only by stimulating DNA-PK but also via P-TEFb activation. Jurkat-pHR’P-Luc cells were treated with increasing doses of cocaine (5, 10, 15, 20, and 25 µM) for 3 h (**A**). Jurkat-pHR’P-Luc cells were treated as follows: untreated and uninfected (Lane 1), infected with HIV (93/TH/051) without cocaine treatment (Lane 2), treated with cocaine without HIV nfection (Lane 3), or pre-treated with different concentrations of cocaine before HIV infection (Lanes 4 to 6) (**C**). Cells were harvested, and nuclear lysates were analyzed by immunoblotting with specific antibodies against P-TEFb subunits CDK9 and Cyclin T1, as well as CDK7 (TFIIH). Actin was used as a loading control. Densitometric analysis of protein bands (normalized to actin) confirmed a significant increase in CDK7, Cyclin T1, and p-CDK9 (Thr186) compared to untreated (Ctrl) cells (**B & D**). Wild type (WT) and DNA-PK knockdown (DNA-PK KD) cells were treated with cocaine for 30 min and 3 h, and nuclear extracts were subjected to mmunoblotting (**E**). Densitometric analysis of protein bands (normalized to actin) showed increased p-CDK9 phosphorylation and CDK7 activation in WT cells upon cocaine exposure (**F**). However, in DNA-PK KD cells, he lack of p-CDK9 (Thr186) phosphorylation and CDK7 activation upon cocaine treatment demonstrated that cocaine-induced activations are DNA-PK specific (**F**). Immunoblots are representative of at least three ndependent experiments. The results are expressed as mean ± SD for three independent experiments, analyzed by two-way ANOVA followed by Tukey’s multiple comparisons test. Asterisks over the bars indicate significant differences: ∗p < 0.05 compared to untreated cells (Ctrl).

To further substantiate that cocaine-induced phosphorylation of CDK9 and activation of CDK7 are due to DNA-PK activation, we conducted experiments using DNA-PK knockdown cells. The wild type (WT) and DNA-PK knockdown (DNA-PK KD) cells were treated with cocaine for 3 h. Subsequently, we analyzed phosphorylation of CDK9 and activation of CDK7. We found a significant reduction to p-CDK9 (Thr186) levels, as well as in total CDK9 and CDK7 following DNA- PK depletion (**Figures 7E & 7F**). Our findings in wild type cells confirmed our above results, indicating that cocaine exposure led to an increase in pCDK9 phosphorylation and activation of CDK7. However, in DNA-PK KD cells, we observed a persistent reduction in pCDK9 (thr186) phosphorylation and CDK7 activation upon cocaine exposure, suggesting that cocaine-induced CDK9 phosphorylation and activation of CDK7 are DNA-PK specific. Together, these findings confirmed our hypothesis by validating that cocaine-induced DNA-PK facilitates the initiation and elongation phases of HIV transcription by stimulating CDK7 (THIIH) and CDK9 (P-TEFb), respectively.

### 3.7. Cocaine-induced DNA-PK relieves the RNAP II pausing by phosphorylating TRIM28 at S824

Later, we examined the impact of cocaine-induced TRIM28 activation (p-TRIM28-(S824) in relieving RNAP II pausing. TRIM28 is one of the RNAP II pausing factors, which restricts the flow of RNAP II on DNA template after transcription of the first 50 to 60 nucleotides. Additionally, it has been recently documented that TRIM28 potently suppresses HIV expression by utilizing both SUMO E3 ligase activity and epigenetic adaptor function [63]. However, phosphorylation of TRIM28 at its Ser824 converts TRIM28 from a pausing factor to transcription-supporting factor [39, 59, 60]. To further extend on our previous findings [39], in this investigation, for the first time, we provide the evidence that DNA-PK is the main kinase that catalyzes the phosphorylation of TRIM28 at Ser824 (p-TRIM28-(S824) and reverses the inhibitory effect of TRIM28 on gene transcription. We hypothesized that if cocaine stimulates DNA-PK and plays a major role in supporting not only initiation but also the elongation phase of HIV transcription, then cocaine-induced DNA-PK should be able to relieve RNAP II pausing, a prerequisite for the elongation phase of transcription. To test this hypothesis, we examined the neutralization of RNAP II pausing through the conversion of TRIM28 from a transcriptionally repressive factor (TRIM28) to a transcriptionally active factor (p-TRIM28 S824) by cocaine-induced DNA-PK-mediated phosphorylation of TRIM28. The THP-1 cells were treated with increasing doses of cocaine for 3 h. The nuclear lysates were analyzed by immunoblotting to detect the phosphorylated form of TRIM28 (p-TRIM28-(S824) and total TRIM28. The expression of Actin protein among samples was evaluated as loading control. As expected, following cocaine exposure, we found enhanced TRIM28 phosphorylation at the position S824 (p-TRIM28-(S824) in a dose-dependent manner (**Figures 8A & 8B**). The densitometric analyses of protein bands further establish the significant dose-dependent increase to p-TRIM28-(S824) levels upon cocaine treatment compared to the untreated cell control. Thus, showing that cocaine by enhancing phosphorylation of TRIM28 relieves the RNAP II pausing during HIV transcription. These results were further confirmed in Jurkat cells (**Figures 8C & 8D**).

**Figure 8:**
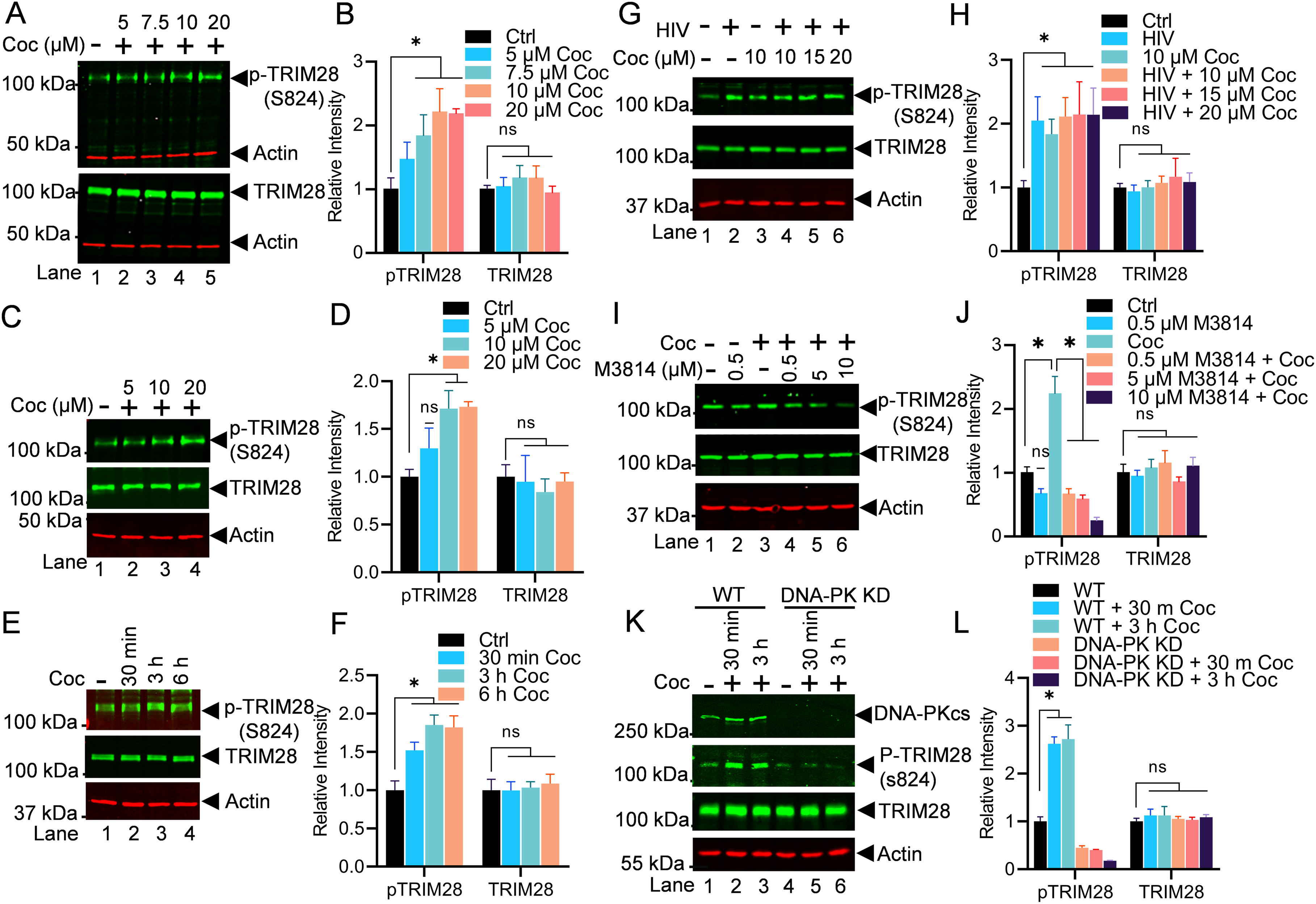
Cocaine-induced DNA-PK relieves RNAP II pausing by phosphorylating TRIM28 at S824. THP-1 (**A & B**) and Jurkat cells (**C & D**) were treated with increasing doses of cocaine, and the nuclear ysates were analyzed via immunoblotting using specific antibodies against pTRIM28 (S824) and total TRIM28. Densitometric analysis confirmed a significant increase in pTRIM28 (S824) levels compared to untreated cells (Ctrl) (**A, B, C & D**). Jurkat-pHR’P-Luc cells were treated with cocaine (10 µM) for varying durations (30 min, 3 h, and 6 h), and the nuclear lysates were analyzed via immunoblotting using specific antibodies against pTRIM28 (S824) and total TRIM28. Densitometric analysis of protein bands (normalized to actin) confirmed a significant increase in pTRIM28 (S824) levels compared to untreated cells (Ctrl) (**E & F**). THP-1 cells were treated as follows: untreated and uninfected (Lane 1), infected with HIV (93/TH/051) without cocaine (Lane 2), treated with cocaine without HIV infection (Lane 3), or pre-treated with different concentrations of cocaine before HIV infection (Lanes 4 to 6). Nuclear lysates were analyzed via mmunoblotting using specific antibodies against pTRIM28 (S824) and total TRIM28 (**G**). Densitometric analysis of protein bands (normalized to actin) confirmed a significant increase in pTRIM28 (S824) levels compared to untreated cells (Ctrl) (**H**). THP-1 cells were treated with different concentrations of M3814 in the presence and absence of cocaine (10 µM), and the nuclear lysates were analyzed via immunoblotting using specific antibodies against pTRIM28 (S824) and total TRIM28 (**I**). Densitometric analysis of protein bands (normalized to actin) confirmed a significant increase in pTRIM28 (S824) levels compared to untreated cells (Ctrl). However, the presence of the inhibitor (M3814) severely impaired pTRIM28 (S824) compared to the cocaine-treated sample (**J**). WT and DNA-PK KD cells were treated with cocaine for 30 min and 3 h, and nuclear lysates were subjected to immunoblotting (**K**). Densitometric analysis of protein bands (normalized to actin) confirmed enhanced phosphorylation of p-TRIM28 in WT cells upon cocaine exposure. However, in DNA-PK KD cells, phosphorylated TRIM28 levels remained reduced upon cocaine exposure, confirming that cocaine-induced TRIM28 phosphorylation is DNA-PK specific (**L**). Immunoblots are representative of at least hree independent experiments. The results are expressed as mean ± SD for three independent experiments, analyzed by two-way ANOVA followed by Tukey’s multiple comparisons test. Asterisks over the bars indicate significant differences. ∗p < 0.05 is for the comparison of cocaine-treated samples against untreated (Ctrl) and he comparison of cocaine plus inhibitors treated against cocaine alone-treated samples

For examining the kinetics of TRIM28 phosphorylation upon cocaine exposure, we treated the Jurkat-pHR’P-Luc cells with a fixed dose of cocaine (10 μM) for different durations: 30 min, 3 h, and 6 h (**Figures 8E & 8F**). Then, we analyzed the nuclear lysates to assess the levels of p-TRIM28 (S824) and TRIM28 total; we noted significant phosphorylation of TRIM28 at 3 h upon cocaine exposure. As anticipated, densitometry analyses revealed a significant increase of TRIM28 phosphorylation following cocaine treatment in a unique kinetics (**Figures 8E & 8F**). Together, our data establish that cocaine-mediated enhanced TRIM28 phosphorylation (p-TRIM28-(S824) plays a crucial role in transitioning HIV transcription from pausing to the elongating phase by antagonizing the pausing effect of TRIM28, and thus, relieving RNAP II pausing.

The results were also reproduced in Jurkat cells infected with replication competent virus (93/TH/051). Jurkat-pHR’P-Luc cells were treated with increasing concentrations of cocaine for 3 h before being infected with 93/TH/051. After 3 h, nuclear extracts were examined for p-TRIM28 (S824) and TRIM28 total. The enhanced phosphorylation of TRIM28 at S824 (p-TRIM28-(S824) upon cocaine treatment was confirmed (**Figures 8G & 8H**).

Subsequently, to determine if the cocaine-induced DNA-PK is responsible for TRIM28 phosphorylation (p-TRIM28 S824), we examined the impact of DNA-PK inhibition on TRIM28 phosphorylation. We found a dose-dependent inhibition of TRIM28 phosphorylation and almost complete elimination of TRIM28 phosphorylation (p-TRIM28 S824) in cells treated with 10 µM M3814 (**Figures 8I and 8J**). Together, these findings confirm that cocaine-induced DNA-PK plays a vital role in RNAP II pause release by enhancing TRIM28 phosphorylation at a specific site (p- TRIM28-(S824), which converts TRIM28 from an inhibitory factor to a transactivator (**Figures 8**).

We further confirmed the specific role of cocaine-stimulated DNA-PK in catalyzing phosphorylation of TRIM28 and reversing its inhibitory effect during HIV transcription by performing experiments using DNA-PK knock down (KD) cells. Cells were infected with lentiviral vectors expressing shRNA either against catalytic subunit of DNA-PK (DNA-PKcs) or scrambled shRNA, which do not target any cellular gene. These cells were treated with cocaine for 30 min and 3 h. Later, phosphorylation of p-TRIM28 at S824 and total TRIM28 was analyzed. In DNA-PK knockdown cells, we observed a clear reduction in the levels of p-TRIM28-(S824) but not TRIM28 (**Figures 8K & 8L**). However, in cells harboring scrambled shRNA, which express normal levels of DNA-PK, we noted enhanced phosphorylation level of p-TRIM28 upon the cocaine exposure, validating our previous results. We also noted the level of phosphorylated TRIM28 remains reduced in the DNA-PK KD cells upon exposure to cocaine, confirming that cocaine induces TRIM28 phosphorylation is DNA-PK specific. Thus, the results demonstrated that the enhanced phosphorylation of TRIM28 induced by cocaine is directly associated with the stimulation of DNA-PK triggered by cocaine (**Figure 8K**).

To understand the cellular kinetics of TRIM28, we analyzed the cytosolic and nuclear levels of p- TRIM28 (S824) and TRIM28 upon cocaine exposure. We also analyzed the impact of cocaine on two main RNAP II pausing factors, namely DSIF (SPT-5) and NELF (NELF-E). Interestingly, we did not observe any significant changes in DSIF and NELF upon cocaine exposure (**Figure 9A, 9B & 9C**). These results clearly document that cocaine primarily relieves RNAP II pausing by inducing DNA-PK mediated phosphorylation of TRIM28 (p-TRIM28-(S824). Altogether, our data validate that cocaine-stimulated DNA-PK relives RNAP II pausing by antagonizing the effect of negative/pausing factors, mainly TRIM28, via its phosphorylation at ser824 (p-TRIM28-(S824), during HIV transcription.

**Figure 9:**
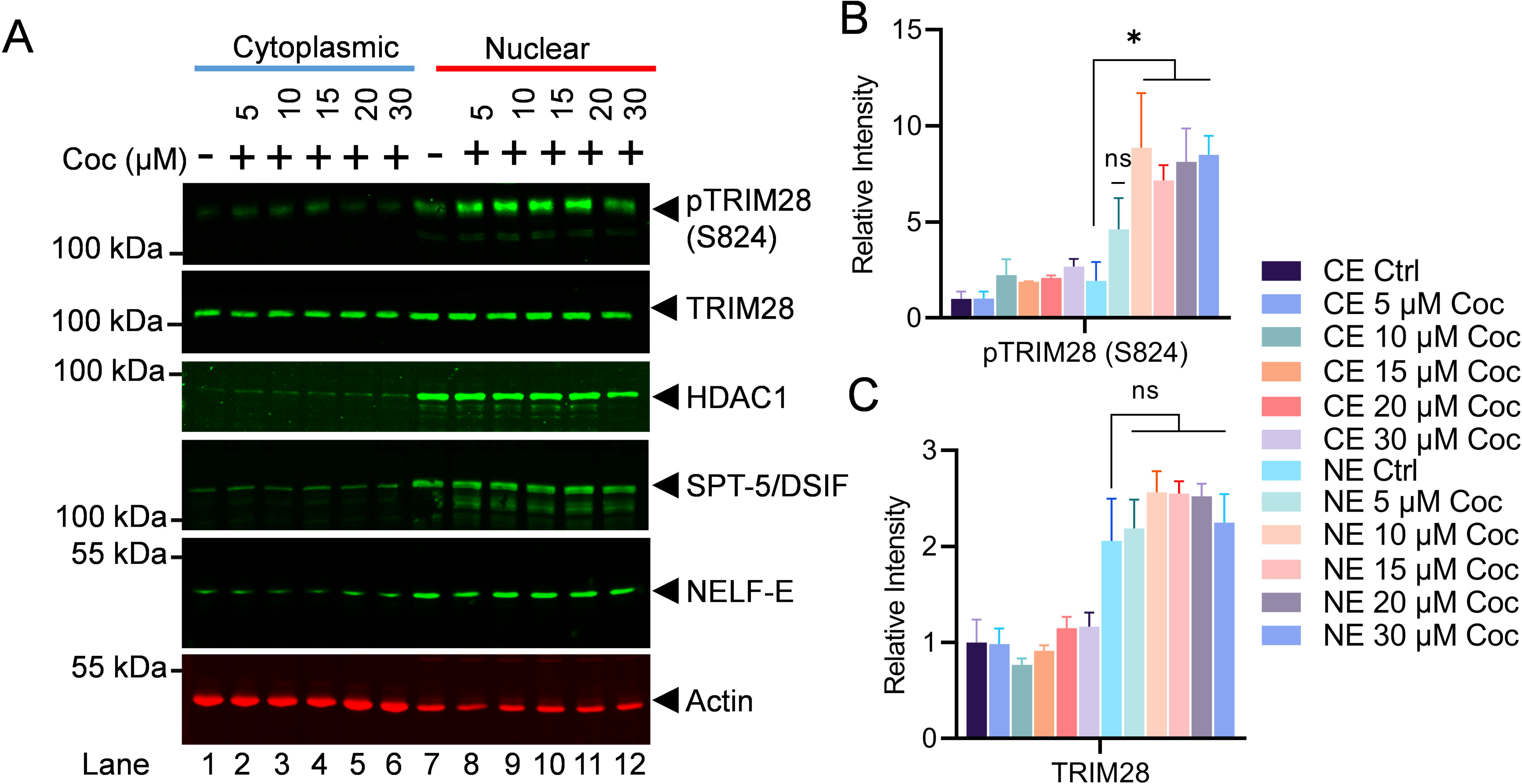
Cocaine promotes RNAP II pause-release by phosphorylating TRIM28 at S824. Jurkat cells were exposed to increasing concentrations of cocaine, and both cytoplasmic and nuclear extracts were subjected to immunoblotting using specific antibodies against pTRIM28 (S824), total TRIM28, DSIF (SPT-5), NELF-E, and HDAC1. Densitometric analysis of protein bands (normalized to actin) confirmed a significant ncrease in nuclear pTRIM28 (S824) levels following cocaine exposure compared to untreated cells (Ctrl) (**A, B & C**). Immunoblots are representative of at least three independent experiments. The results are expressed as mean ± SD for three independent experiments, analyzed by two-way ANOVA followed by Tukey’s multiple comparisons test. Asterisks over the bars indicate significant differences. Statistical significance is set as p < 0.05 (∗) compared to untreated cells (Ctrl).

### 3.8. Cocaine boosts HIV transcription by enhancing the recruitment of DNA-PK and pTRIM28 at HIV LTR promoter

Previously, we documented the parallel presence of DNA-PK along with RNAP II throughout the HIV proviral genome during HIV transcription [37, 39]. Additionally, we have shown the recruitment of TRIM28 at HIV long terminal repeat (LTR) during HIV transcription [39]. We also found that cell activation enhances both the nuclear translocation and LTR recruitment of DNA-PK [39]. Given that cocaine further augments the nuclear translocation of DNA-PK, we hypothesize that enhanced nuclear translocation of DNA-PK should translate into higher recruitment of DNA-PK and TRIM28 at HIV LTR. To test this hypothesis, we evaluated the recruitment of DNA-PK and p-TRIM28-(S824) at HIV LTR in the presence and absence of cocaine by chromatin Immunoprecipitation (ChIP) assay using our standard methodology [28, 37, 68, 93]. The ChIP assays were performed using antibodies, namely IgG (control), DNA-PKcs, RNAP II, CDK7, CDK9, pTRIM28 (S824), and H3K27me3. The analysis was done in Jurkat cells freshly infected with p-HR’P-Luc (**Figure 2A**). The recruitment of RNAP II at HIV LTR was assessed as positive control to mark ongoing HIV transcription. We examined CDK7 as a marker of transcriptional initiation, as CDK7 (TFIIH) plays a role during the initiation phase of HIV transcription. The recruitment of CDK9 (P-TEFb) at HIV LTR was evaluated to indicate the elongation phase of HIV transcription, as recruitment of P-TEFb is crucial to support HIV transcriptional elongation. The immunoprecipitated DNA was analyzed using four primer sets targeting different regions of HIV LTR. The first primer set amplifies the promoter region of the LTR (-116 to +4 with respect to the transcription start site, **Figures 10A & 10E**). The second primer set amplifies the Nuc-1 region of the LTR (+30 to +134 with respect to the transcription start site, **Figures 10B & 10F**). The factors that mainly bind at the promoter and Nuc-1 region usually mark factors involved in the initiation phase of HIV transcription. The third primer set amplifies the downstream Nuc-2 region of the LTR (+283 to +390 with respect to the transcription start site, **Figure 10C & 10G**). The fourth primer set amplifies further downstream ENV region of HIV (+2599 to +2697, **Figure 10D & 10H**). The factors that bind around Nuc-2 region and downstream primarily represent those involved in the elongation phase of transcription. Following cocaine treatment, as anticipated, we found higher recruitment of RNAP II showing upregulation of HIV transcription. Moreover, enhanced RNAP II levels at promoter, Nuc-1, Nuc-2, and Env region of provirus in cocaine treated cells indicate enhanced ongoing HIV gene expression upon cocaine treatment. Interestingly, in parallel to the recruitment of RNAP II, we observed significantly enhanced recruitment of DNA-PKcs at the promoter, Nuc-1, Nuc-2, and the Env regions of LTR following cocaine treatment (**Figures 10A, 10B, 10C & 10D**). These results corroborate our previous data, where we showed the continuous presence and gliding of DNA-PKcs with RNAP II along the HIV genome during transcription [37]. Notably, we also found enrichment of p-TRIM28- (S824) at the promoter and Nuc-1 region (**Figures 10E & 10F**). However, we did not observe significant changes in the Nuc-2 and Env region of HIV LTR (**Figures 10G & 10H**). Meanwhile, we noted substantially higher recruitment of CDK7 (kinase subunit of TFIIH) at promoter and Nuc-1 of the LTR region. However, we observed a decrease of CDK7 recruitment at Nuc-2 but no significant changes in the Env region (**Figures 10E, 10F, 10G & 10H**). The finding that after cocaine stimulation, CDK7 was enriched at the LTR promoter but not at the Nuc-2 region validates its involvement specifically during the initiation phase of HIV transcription. Interestingly, the loss of H3K27Me3 from HIV LTR following cocaine treatment demonstrates the removal of transcriptionally repressive (heterochromatin) structure and establishment of transcriptionally active (euchromatin) structure at HIV LTR following cocaine treatment. These data further validate our previous findings, where we showed that cocaine enhances HIV transcription by promoting euchromatin structure at HIV LTR [28]. As anticipated, following cocaine exposure, we also found enhanced recruitment of CDK9 (kinase subunit of P-TEFB) specifically at the downstream region of LTR but not much at the promoter region, validating its role during the elongation phase of transcription (**Figure 10G & 10H**). Following cocaine exposure, the specific enrichment of CDK9 (P-TEFb) at the downstream region of LTR and CDK7 (TFIIH) at the promoter region validates the authenticity of our assay system and ChIP analysis.

**Figure 10:**
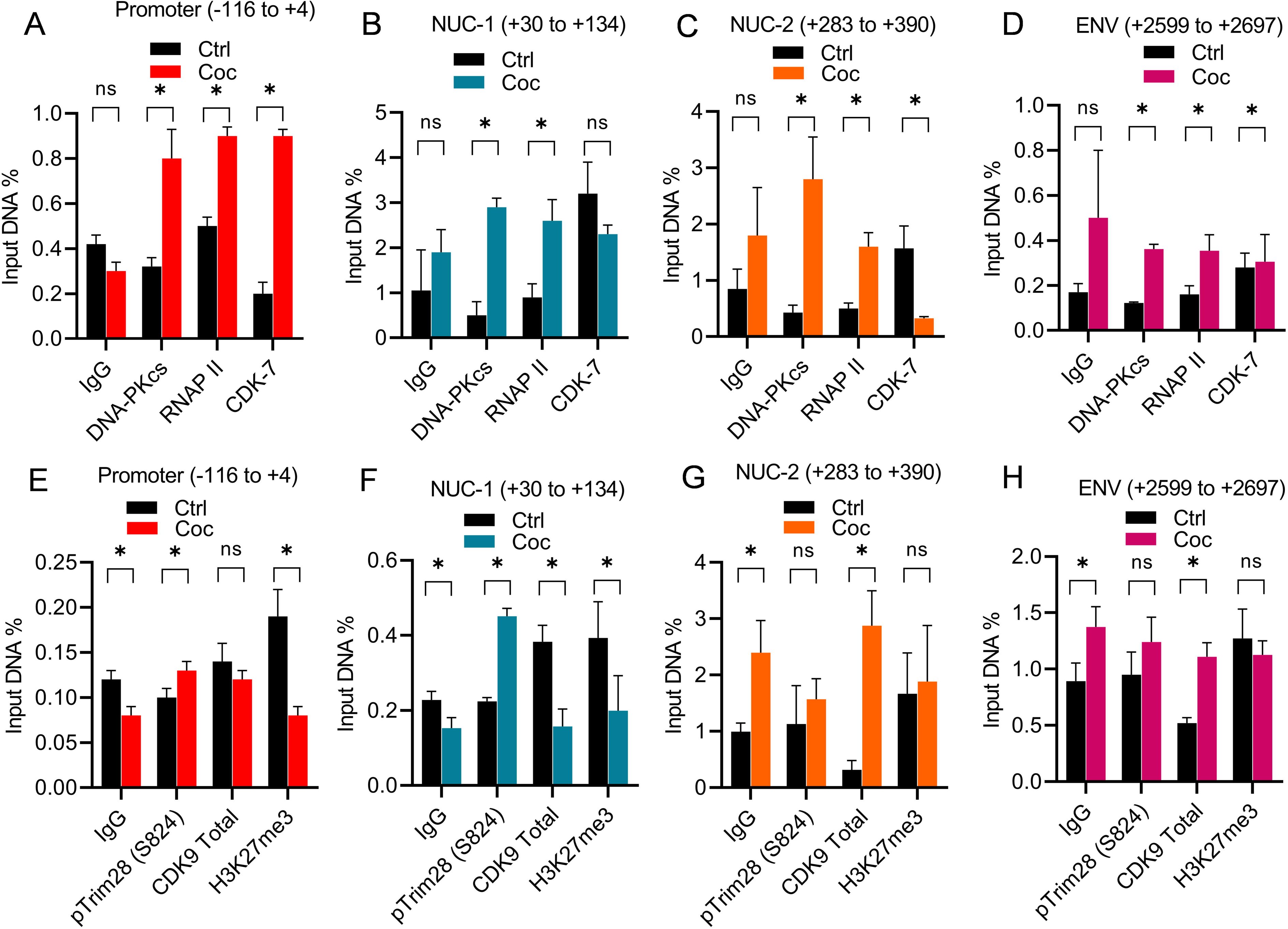
Cocaine enhances HIV transcription by promoting recruitment of DNA-PKcs and pTRIM28 (S824) at HIV LTR promoter. Jurkat cells freshly infected with replication-incompetent HIV, pHR’P-Luc, were exposed to cocaine. Chromatin immunoprecipitation (ChIP) assays were conducted to assess the recruitment kinetics of DNA-PKcs, RNAP II, CDK7 (TFIIH), pTRIM28 (S824), total CDK9, and H3K27me3 at the promoter (**A & E**), Nucleosome-1 (**B & F**), Nucleosome-2 (**C & G**), and downstream Envelope regions (**D & H**) of HIV LTR, using specific primer sets. The results are presented as mean ± SD for three independent experiments, analyzed by two-way ANOVA followed by Tukey’s multiple comparisons test. Asterisks above the bars ndicate significant differences. Statistical significance is set as p < 0.05 (∗) compared to untreated cells (Ctrl).

Overall, our results demonstrate that cocaine stimulates and enhances the nuclear translocation and catalytic activity of DNA-PK (p-DNA-PK S2056), which leads to its higher recruitment at HIV LTR. DNA-PK subsequently catalyzes the phosphorylation of TRIM28 (p- TRIM28 S824) and converts TRIM28 from a pausing factor to a transcription activator. Overall, these modifications relieve RNAP II pausing and promote HIV transcriptional elongation, a necessity to make complete HIV genomic transcripts, which are required for generating viral progeny.

### 3.9. Cocaine induced DNA-PK activation promotes HIV transcription by supporting several aspects of HIV transcription

To summarize our findings from current and previous investigations, we present the following model for DNA-PK role during HIV transcription (**Figure 11**) [37, 39]. In our previous studies, we have established the association of DNA-PK and RNAP II along HIV proviral DNA template throughout HIV gene expression. In this study, we found that cocaine exposure augments the nuclear translocation and functional activation of DNA-PK (p-DNA-PK S2056). DNA-PK subsequently facilitates the multiple critical phases of HIV transcription, namely initiation, RNAP II pause release, and elongation. Cocaine-induced DNA-PK promotes the initiation phase of transcription by catalyzing the phosphorylation of RNAP II CTD at Ser5. In addition, cocaine-stimulated DNA-PK facilitates the elongation phase of HIV transcription by both directly catalyzing and promoting the recruitment of P-TEFb for the phosphorylation of Ser2 within the RNAP II CTD. The hyperphosphorylation of RNAP II CTD at Ser2 makes RNAP II processive or elongation proficient. Another noteworthy finding is that cocaine-stimulated DNA-PK relieves the RNAP II pausing selectively through TRIM28 by catalyzing TRIM28 phosphorylation at Ser824 (p-TRIM28 S824). This modification transforms TRIM28 from a transcription pausing factor to a transcription- supporting factor. Thus, phosphorylation of TRIM28 at Ser824 relieves RNAP II pausing and allows RNAP II to proceed along DNA template or transcriptional elongation. Our findings collectively underscore the profound impact of cocaine-induced DNA-PK activation on various facets of HIV transcription, ultimately culminating in the potent promotion of viral gene expression. Therefore, DNA-PK inhibitors profoundly inhibit HIV transcription, replication, and latency-reactivation.

**Figure 11:**
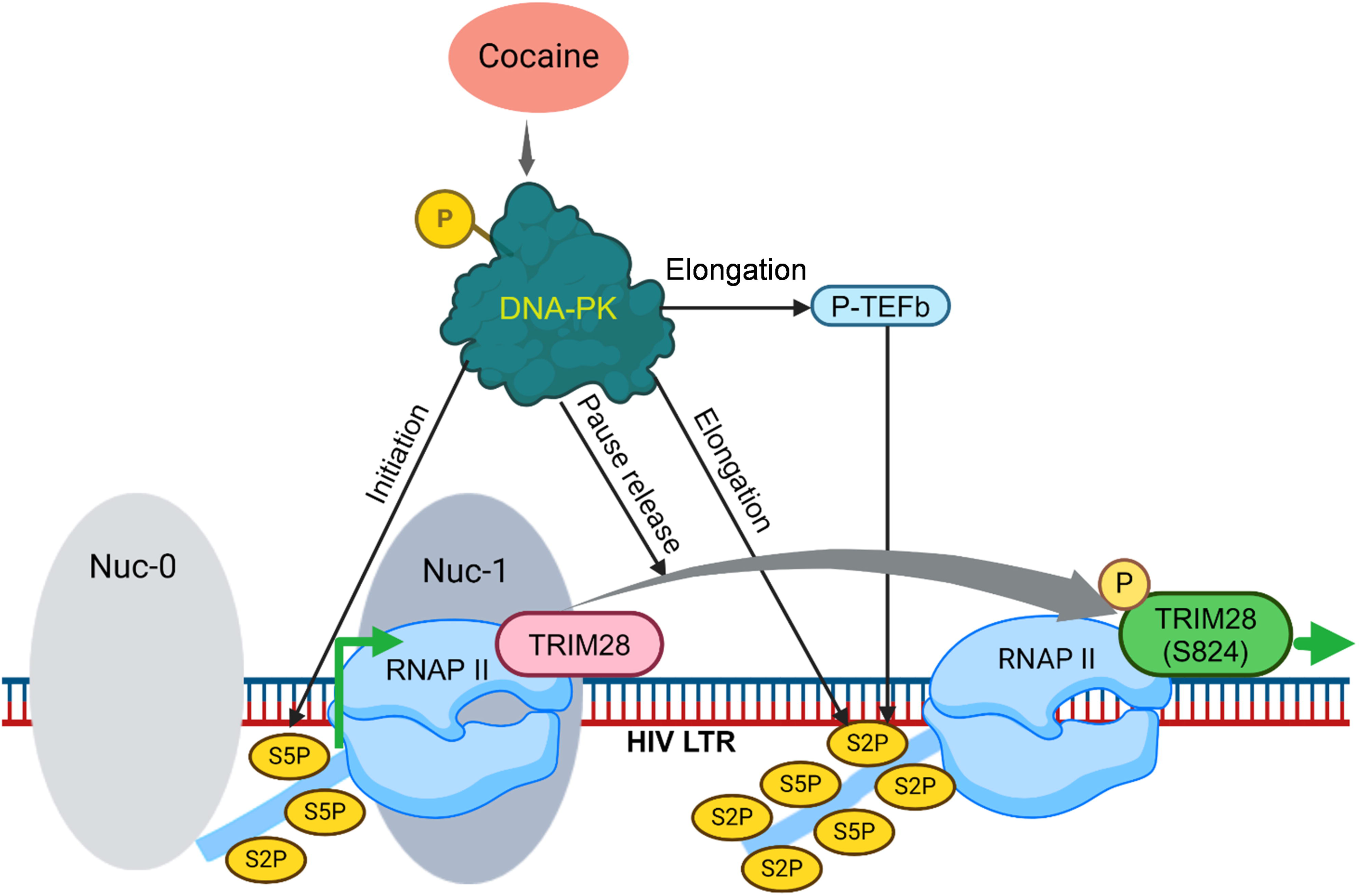
Cocaine-mediated DNA-PK activation enhances multiple aspects of HIV transcription. Cocaine-induced DNA-PK activation facilitates various stages of HIV transcription. Firstly, it enhances the nitiation phase by phosphorylating the C-terminal domain (CTD) of RNA polymerase II (RNAP II) at Ser5. Secondly, cocaine-stimulated DNA-PK promotes the elongation phase by both directly catalyzing and acilitating the recruitment of positive transcription elongation factor b (P-TEFb), leading to the phosphorylation of Ser2 within the RNAP II CTD. This posttranslational modification renders RNAP II processive, ensuring efficient elongation. Finally, cocaine-induced DNA-PK activity also alleviates RNAP II pausing by phosphorylating TRIM28 at Ser824 (p-TRIM28 S824). This modification transforms TRIM28 from a transcriptional pausing factor to a facilitator (transactivator), thereby supporting HIV transcription.

## 4. Discussion

HIV/AIDS remain a dreadful disease, as an effective vaccine or cure is yet to develop [5, 94–98]. Nevertheless, with the introduction of ART, the quality of PLWH significantly increases [1, 6]. However, one has to rely on medication for the rest of one’s life to keep control of HIV disease progression. The anti-HIV therapy, ART, is highly effective in suppressing viral replication, maintaining healthy immune system, and reducing risk of HIV transmission. Unfortunately, cocaine, one of the most abused drugs by HIV patients, can disrupt regular activities potentially leading to inconsistent or missed doses of ART. Poor adherence usually leads to treatment failure, development of drug resistant HIV strain and compromised immune functions [99, 100]. Cocaine further affects the normal functioning of immune cells, suppressing the immune system and exacerbating the effect of HIV infection leading to faster disease progression, specially making HIV patients vulnerable to opportunistic infections. Furthermore, given that cocaine strongly impacts brain functioning, cocaine use by HIV patients not only accelerates HIV replication in the CNS but also exacerbates normal brain functioning. The interaction between cocaine and HIV is a multifaceted and concerning issue. Therefore, understanding the molecular mechanisms that govern HIV life cycle, especially transcription and replication, are crucial for relieving from HIV and cocaine induced neurotoxicity in addition to HIV cure and eradication [16, 17, 27, 39]. In this study, we showed the pivotal role played by cocaine-induced activation of DNA-PK in bolstering various stages of HIV transcription, consequently augmenting HIV replication. Our investigation has unveiled that cocaine significantly upregulates the expression of DNA-PK, prompts its translocation into the nucleus, and enhances the functional activity of DNA-PK by enhancing its phosphorylation at S2056. Subsequently, the cocaine-induced DNA-PK facilitates transcriptional initiation by augmenting the phosphorylation of CTD at Ser5, relieves RNAP II pausing through TRIM28 phosphorylation at S824, and promotes transcriptional elongation both by directly catalyzing the phosphorylation of CTD at Ser2 and through P-TEFb stimulation and recruitment. Accordingly, upon specific inhibition or depletion of DNA-PK using specific inhibitors or knockdown, respectively, we found profound restriction to cocaine-induced HIV transcription and replication. These collective results unveil the underlying molecular mechanisms through which cocaine-induced DNA-PK stimulation augments HIV transcription and replication.

DNA-PK is a serine/threonine protein kinase complex composed of a heterodimer of Ku proteins (Ku70/Ku80) and a catalytic subunit DNA-PKcs [40, 41]. DNA-PK is a critical component of the cellular response following DNA damage [40, 41]. DNA-PK is one of the main components of DNA repair pathway upon double-strand breaks, especially in the NHEJ DNA double-strand break repair pathway [43, 44]. Therefore, DNA-PK is extensively studied in DNA double strand break repair. The DNA-PK role in HIV transcription was first identified as a complex that phosphorylates the transcription factor SP1 [101] and as a interacting component of RNAP II [47]. Nevertheless, its role in transcription was understudied. For the first time, we demonstrated the precise role of DNA-PK during any transcription process by defining the mechanism through which DNA-PK promotes HIV transcription and involved mechanisms [37, 39]. Later, several studies emerged that further strengthened the link between DNA-PK and transcriptional regulation [102]. Given that, HIV transcription is the fundamental step that plays a crucial role in regulating HIV replication and latency-reactivation. In our previous studies we have documented the underlying molecular mechanism through which DNA-PK promotes HIV transcription [39]. Moreover, we found that cocaine also enhances HIV transcription and replication [16, 27, 28]. These facts prompted us to study if the cocaine-enhanced HIV transcription and replication is due to the activation of DNA-PK. In this investigation, we demonstrated that cocaine significantly upregulates nuclear level of DNA- PK and augments its activity by enhancing its phosphorylation at Serine 2056 residues. We reproduced these findings in cells of different lineages, including both lymphoid and myeloid lineages. Given that in our previous findings we noted higher recruitment of DNA-PK at HIV LTR following cell stimulation [37, 39], we evaluated if cell stimulation by cocaine also results in enhanced nuclear translocation of DNA-PK. We found that cocaine-induced cell stimulation was sufficient and promoted the nuclear translocation of DNA-PK (**Figure 1**). Interestingly, the nuclear levels of the DNA-PK significantly increased following cocaine treatment with the corresponding decrease in the cytoplasmic levels, indicating the translocation of DNA-PK towards the nucleus (**Figures 1I & 1J)**. Subsequently, we analyzed the corresponding upregulation of DNA-PK recruitment of DNA-PK due to higher availability of DNA-PK in the nucleus by ChIP assay. As expected, upon cocaine exposure, we found a notable increase in the recruitment of DNA-PK. Additionally, along with DNA-PK, we found the corresponding higher recruitment of RNAP II at HIV LTR following cocaine treatment (**Figure 10A, 10B, 10C & 10D**). This finding reaffirmed our prior findings where we established DNA-PK interaction with RNAP II and showed parallel-enhanced recruitment of both DNA-PK and RNAP II following cell stimulation [37]. Interestingly, paralleling the recruitment of RNAP II, we also noted an augmented recruitment of DNA-PK not only at the promoter and Nuc-1 regions but also at the downstream regions of the HIV genome (**Figure 10A, 10B, 10C & 10D**). This validates the role of DNA-PK in different phases of HIV transcription, including initiation, RNAP II pause release, and elongation phases. Accordingly, we found higher levels of RNAP II at the promoter, Nuc-1, and Env region of the provirus, signifying enhanced ongoing HIV gene expression following cocaine exposure. This observation further strengthens our previous results proposing that DNA-PK and RNAP II are part of a larger transcription complex [37, 39]. Later, we assessed if HIV infection promotes cell stimulation and consequently DNA-PK activation. Notably, we found significant upregulation of DNA-PK and its activation (p-DNA-PK S2056), suggesting crucial role of DNA-PK during HIV transcription. Together, these findings underscore the intricate relationship between cocaine exposure and the LTR recruitment of DNA-PK, shedding light on the potential mechanism through which cocaine augments HIV transcription.

Our previous findings, where we establish the vital role of DNA-PK during HIV transcription [37, 39], has been extended by others. The role of DNA-PK in general cell transcription has also been documented [60], validating the important role of DNA-PK during basic transcriptional process. To further validate our findings and establish the crucial role of cocaine in stimulating DNA-PK during HIV transcription, we employed a highly specific DNA-PK inhibitor, M3814. The dose-dependent inhibition of HIV transcription by M3814, indicated by reduced luciferase gene expression from LTR promoter (**Figure 3B & 3C**), confirmed the direct role of DNA-PK in promoting HIV transcription. Given that, TNF-α was unable to reactivate the latent HIV in the presence of M3814, demonstrating that DNA-PK inhibitors could be useful in restricting the reactivation of latent HIV provirus as well (**Figure 3E & 3F**). Interestingly, we did not observe any noticeable cell toxicity with the used concentrations of M3814 (**Figure 3D**), establishing the physiological significance of the findings. Subsequently, we also evaluated the effect of two different DNA-PKcs inhibitors, M3814, and NU7441 on HIV replication. We found that the more specific DNA-PK inhibitors (DNA-PKi) were better at repressing HIV gene expression and replication (**Figure 4**). This observation again confirmed the target-specific impact of DNA-PKi. Moreover, cell viability analysis validated the physiological viability of the pre-clinically and clinically tested DNA-PK inhibitors as potential HIV therapeutics.

Previously, we identified both the presence of DNA-PK at HIV LTR and direct catalyzation of RNAP II CTD phosphorylation by DNA-PK [37, 39]. We investigated whether cocaine induced HIV transcription and replication is also due to DNA-PK stimulation and subsequently RNAP II CTD phosphorylation, we examined the state of RNAP II CTD phosphorylation following cocaine exposure. The significant upregulation of Ser2 and Ser5 phosphorylation following cocaine treatment in a dose-dependent manner confirmed that cocaine augments HIV transcription by supporting RNAP II CTD phosphorylation (**Figure 6A & 6B**). Given that, Ser5 phosphorylation is the marker of transcriptional initiation, and Ser2 phosphorylation is linked to the elongation phase of transcription, including HIV transcription. The data obtained showed that cocaine facilitates both the initiation and elongation phases of transcription. The results were reproduced in the cells of multiple lineages to show the ubiquitous prevalence of the observed phenomenon (**Figure 6C, 6D, 6E & 6F**). Subsequently, we explored whether cocaine-enhanced RNAP II phosphorylation is a result of DNA-PK activation using a clinically evaluated highly specific DNA-PK inhibitor (M3814) in the presence of cocaine (**Figures 6E & 6F)**. The dose-dependent inhibition of RNAP II CTD phosphorylation at both Ser2 and Ser5 sites by M3814 validated the specific role of DNA-PK in catalyzing CTD phosphorylation. Altogether, our findings confirmed our hypothesis that cocaine, through activation of DNA-PK, significantly influences both the initiation and elongation phases of HIV transcription, contributing to a more comprehensive understanding of the molecular mechanism behind cocaine’s impact on HIV gene expression.

The previous studies have established the CDK9 subunit of P-TEFb as the main player that promotes RNAP II processivity by catalyzing RNAP II CTD phosphorylation at Ser2 position; thus, supports the elongation phase of transcription [103, 104]. Therefore, we sought to investigate the nuclear level of P-TEFb. Analyzing the nuclear level, our results suggested that cocaine significantly enhances the phosphorylation of CDK9 and Cyclin T1, indicating that cocaine further supports the ongoing elongation phase of HIV transcription through P-TEFb stimulation. Nevertheless, cocaine does not affect the Total CDK9 level. Later, we examined the impact of cocaine on the initiation phase of HIV transcription, and, as anticipated, we found significant upregulation of CDK7, a subunit of TFIIH that is well known to support the initiation phase of transcription, including HIV transcription. These findings were validated in different cell types, both myeloid and lymphoid cells (**Figures 7A, 7B, 7C, 7D, 7E & 7F**). To further validate that cocaine-induced phosphorylation of CDK9 and activation of total CDK7 are indeed reliant on the specific activation of DNA-PKcs, we conducted experiments using a DNA-PKcs knockdown cell line exposed to cocaine. In the absence of DNA-PKcs, we observed a marked decrease in p-CDK9 (Thr186) levels, as well as a reduction in total CDK9 and CDK7. Notably, in wild-type cells, exposure to cocaine resulted in the anticipated enhancement of CDK7 and CDK9 phosphorylation, consistent with our previous findings. However, in DNA-PKcs knockdown cells, the levels of pCDK9 (Thr186) and CDK7 remained reduced following cocaine exposure, providing strong evidence that cocaine-induced CDK9 phosphorylation and CDK7 activation are specifically mediated by DNA-PKcs. The impact of cocaine on both the initiation and elongation phases of HIV transcription was further validated by showing the presence of TFIIH (CDK7) and P-TEFb (CDK9 and CyclinT1), respectively, at HIV LTR (**Figure 10**) through ChIP assays, upon cocaine treatment. Given that P-TEFb plays a crucial role during the elongation phase, accordingly we found specific enrichment of CDK9 at the downstream region, namely, Nuc-2 and Env region of HIV, but highly reduced recruitment at promoter and the Nuc-1 region in cocaine treated cells (**Figure 10**). Similarly, after cocaine stimulation, CDK7 was enriched as expected at the LTR promoter but not at the Nuc-2 region, again validating its requirement especially during the initiation phase of HIV transcription. Our previous findings showed not only the direct interaction between DNA-PK and RNAP II, but also parallel recruitment of DNA-PK along RNAP II at HIV LTR upon cell stimulation [37, 39]. In addition, we have shown cell stimulation following cocaine exposure [27]. These findings prompted us to investigate whether cocaine-mediated cell stimulation and induced DNA-PK activation enhances RNAP II CTD phosphorylation, both via directly catalyzing and through promoting P-TEFb recruitment at HIV LTR. As expected, we found parallel recruitment of DNA-PK and RNAP II along HIV genome following cocaine treatment, confirming that cocaine-induced cell stimulation is sufficient not only to activate DNA-PK (**Figure 1**), but also to enrich DNA-PK at HIV LTR proportional to RNAP II recruitment at LTR (**Figure 10**). Interestingly, the decrease in the recruitment of H3K27Me3 at HIV LTR following cocaine treatment demonstrates the loss of repressive epigenetic structure, and establishment of transcription-supporting euchromatin structure, aligning with our previous findings [39]. Altogether, these findings demonstrate that cocaine-induced DNA-PK facilitates transcriptional initiation by catalyzing the RNAP II CTD at Ser5. Furthermore, cocaine-mediated DNA-PK stimulation augments the elongation phase of HIV transcription by enhancing the phosphorylation of RNAP II CTD at Ser2 both via directly catalyzing and promoting the recruitment of P-TEFb.

We also explored whether cocaine can facilitate HIV transcription by promoting RNAP II pause release. We found that cocaine profoundly enhances TRIM28 phosphorylation at its serine 824 residue. This specific phosphorylation event relieves the TRIM28-mediated pausing to RNAP II and even converts TRIM28 into a transcription-supporting factor [58, 60]. The established interaction between TRIM28 and RNAP II underscores the significant role of TRIM28 in regulating HIV transcription. Additionally, our studies, in line with previous research, have elucidated that DNA-PK interacts with TRIM28 and catalyzes its phosphorylation at serine 824, resulting in the formation of p-TRIM28-(S824) [39, 60]. This phosphorylation event has been associated with positive elongation factors, suggesting its potential role in facilitating the transition from transcriptional pausing to elongation. Consequently, this modification transforms TRIM28 from a transcriptionally repressive factor into a transcriptionally active one. Therefore, we investigated whether cocaine can convert TRIM28 from a transcriptionally repressive factor to a transcriptionally active one by examining the phosphorylation of TRIM28 at S824. We observed that, upon cocaine exposure, the phosphorylation of TRIM28 at S824 significantly increases in a dose dependent manner (**Figure 8A & 8B**). These findings were confirmed in cells of different lineages, validating the uniformity of the findings (**Figure 8C, 8D, 8E, 8F, 8G, 8H, 8I & 8J**). Later, we analyzed both cytosolic and nuclear levels of p-TRIM28 (S824) and TRIM28 upon cocaine exposure. We noted a significant increase in nuclear levels of p-TRIM28 (S824) in cocaine treated cells, but TRIM28 total did not change significantly (**Figure 9A**). These findings further validated the cocaine-induced activation and phosphorylation of TRIM28 at S824. Subsequently, we analyzed the recruitment of p-TRIM28- (S824) at HIV LTR using ChIP assays. As anticipated, we noted enhanced recruitment of phosphorylated TRIM28 (S824) in parallel to DNA-PK recruitment along HIV genome after cocaine treatment (**Figure 10E & 10F**). The accumulation of p-TRIM28 (S824) marks the presence of the transcription-supporting form of TRIM28 and thus indicates the transformation of paused RNAP II into a processive elongating RNAP II. This observation strongly suggests that by enhancing the phosphorylation of TRIM28, cocaine effectively alleviates RNAP II pausing, thereby providing essential support to the process of HIV transcription. This is another molecular mechanism through which cocaine influences the regulation of transcriptional processes, specifically within the context of HIV gene expression. We further investigated whether cocaine-induced phosphorylation of TRIM28 at S824 is a result of cocaine induced-DNA-PK activation. Upon treating cells with a specific DNA-PK inhibitor, we observed dose-dependent inhibition of cocaine-induced phosphorylation of TRIM28 at S824 (**Figure 8I & 8J**). This finding confirms the critical role played by DNA-PK in promoting RNAP II pause release by selectively catalyzing TRIM28 phosphorylation at S824 and subsequently promoting HIV transcription following cocaine exposure.

Overall, our findings presented here provide compelling and robust evidence affirming the pivotal role played by cocaine on HIV transcription and gene expression. Our investigations have revealed that cocaine significantly upregulates the nuclear levels of DNA-PK, augments its catalytic activity through specific phosphorylation at S2056, besides enhancing its nuclear translocation. We found that cocaine-induced activation of DNA-PK significantly contributes to various stages of HIV transcription, subsequently bolstering the process of HIV replication. Specifically, the activation of cocaine-induced DNA-PK assumes a critical role in facilitating transcriptional initiation by augmenting the phosphorylation of RNAP II CTD at Ser5, alleviating RNAP II pausing through the phosphorylation of TRIM28 at S824 and promoting transcriptional elongation through both the catalysis of CTD phosphorylation at Ser2 and the enhancement of P-TEFb activity. It is noteworthy that our observations have distinctly demonstrated that inhibition or depletion of DNA-PK results in a substantial impediment to cocaine-induced HIV transcription and replication. The overall findings suggest a comprehensive insight into the underlying molecular mechanisms by which cocaine- induced DNA-PK effectively elevates HIV transcription and gene expression (**Figure 11**).

Additionally, we have established the translational potential of DNA-PK inhibitors in curtailing HIV gene expression, replication, and the reactivation of latent provirus. These outcomes advocate for the potential therapeutic application of specific DNA-PK inhibitors as adjuncts in ART regimens, thereby augmenting the efficacy of anti-HIV therapy and potentially curbing the incidence of HIV- associated cancers, given that DNA-PK inhibitors are currently under investigation for cancer treatment. It is noteworthy that while anti-retroviral therapy (ART) treatment effectively controls HIV replication, it is ineffective in regulating HIV gene expression from reactivated latent provirus. These findings strongly advocate for the inclusion of transcriptional inhibitors, such as DNA-PK inhibitors, to supplement ART regimens to mitigate the transient reactivation of latent proviruses, confirmed also by our previous findings involving HIV patients’ samples [39].

## Limitation of the study

Our study has a limitation. We did not consistently quantify the precise amount of replication- competent viruses used for cell infection. However, we maintained an equal viral load in both the control (mock) and test samples.

## Conclusion

Understanding the molecular mechanisms that control the HIV life cycle, particularly in transcription and replication, is crucial for HIV cure and eradication. Our research findings presented herein provide strong and compelling evidence for the important role of cocaine-induced activation of DNA- PK in supporting various phases of HIV transcription, subsequently bolstering HIV replication. Our investigations have revealed that cocaine significantly upregulates the levels of DNA-PK expression, triggers the activation of DNA-PK through enhanced phosphorylation at S2056, and induces its translocation into the nucleus. The activation of cocaine-induced DNA-PK plays a crucial role in promoting transcriptional initiation by enhancing the phosphorylation of CTD at Ser5, alleviating RNAP II pausing by phosphorylating TRIM28 at S824 and facilitating transcriptional elongation by both catalyzing the phosphorylation of CTD at Ser2 and enhancing the P-TEFb recruitment. Notably, our data demonstrate that inhibiting or depleting DNA-PK severely impedes cocaine-induced HIV transcription and replication. These results collectively unveil the underlying molecular mechanisms through which cocaine-induced DNA-PK enhances HIV transcription and gene expression.

## Supporting information

Sharma et. al Supplementary file

## Acknowledgements

We are grateful to the National institute of Health for the research grant to M.T. We are also thankful to the Flow Cytometry core facility of Thomas Jefferson University. We would like to thank Dr. Kartik Krishnamurthy for the support and assistance. We also want to thank the reviewers for providing constructive criticisms prior to publication. MT-4 cells were obtained through the NIH AIDS Reagent Program, Division of AIDS, NIAID, NIH: MT-4 from Dr. Douglas Richman. Moreover, we would like to thank the Center for Translational Medicine, Thomas Jefferson University, including all staff members for their technical support and assistance in conducting the experiments for this study. We express our gratitude to Liz Declan for her assistance in editing the manuscript for English language clarity.

## Funding

M.T. was supported by research grants (Grant Nos.: R01DA041746 and 1R21MH126998-01A1) from the National Institute on Drug Abuse (NIDA) and the National Institute on Mental Health (NIMH) of the National Institute of Health. The funders had no role in study design, data collection, and analysis, decision to publish, or preparation of the manuscript.

## Contributions

The research was conceptualized by MT and planned by ALS and MT. ALS, PT, and MK conducted the experiments. ALS and MT carried out data analysis and prepared the initial manuscript draft. Both ALS and MT contributed to manuscript revisions. MT oversaw the project and secured funding. All authors participated in reviewing and approving the final manuscript version.

## Ethics declarations

Ethics approval and consent to participate: Not applicable.

## Consent for publication

Not applicable

## Competing interests

The authors declare no competing interests.

## Data Availability

The datasets generated from this study are included in this manuscript.

## Key resources table

| REAGENT or RESOURCE | SOURCE | IDENTIFIER |
| --- | --- | --- |
| **Antibodies** |  |  |
| *Anti-HIV1 p55 + p24 + p17 antibody* | Abcam (Cambridge, UK) | Cat# ab63917; RRID: AB_1139524 |
| pDNA-PK S2056 | Cell signaling Technology (Danvers, MA, USA) | Cat# E9J4G |
| DNA-PKcs | Santa Cruz (Dallas, TX, USA) | Cat# sc-390698 |
| Histone Deacetylase 1 (HDAC1) Antibody (10E2): | Santa Cruz (Dallas, TX, USA) | Cat# sc-81598 |
| pTRIM28 (S824) | Bethyl Lab (Montgomery, TX, USA) | Cat# A300-767A-M |
| TRIM28 total | Bethyl Lab (Montgomery, TX, USA) | Cat# A300-275-A-T |
| Pol II (CTD4H8) | Santa Cruz (Dallas, TX, USA) | Cat# sc-47701; RRID: AB_677353 |
| CDK7 (c-4) | Santa Cruz (Dallas, TX, USA) | Cat# sc-7344 |
| CDK9 (C12F7) | Cell signaling Technology (Danvers, MA, USA) | Cat# 2316 |
| NF-kb p-65 | Santa Cruz (Dallas, TX, USA) | Cat# sc-8008 |
| Phospho-CDK9 (Thr186) | Cell signaling Technology (Danvers, MA, USA) | Cat# 2549 |
| Cyclin T1 (D1B6G) | Cell signaling Technology (Danvers, MA, USA) | Cat# 81464 |
| Tri-Methyl-Histone H3 (Lys27) (C36B11) | Cell signaling Technology (Danvers, MA, USA) | Cat# 9733 |
| SPT5 Antibody | Cell signaling Technology (Danvers, MA, USA) | Cat# 9033 |
| NELF-E antibody (F-9) | Santa Cruz (Dallas, TX, USA) | Cat# sc-377052 |
| RNA Pol II Ser2 | Bethyl Lab (Montgomery, TX, USA) | Cat# A300-654A-M; RRID: AB_2779370 |
| RNA Pol II Ser5 | Bethyl Lab (Montgomery, TX, USA) | Cat# A304-408A-M; RRID: AB_2781885 |
| Actin (c4) | Santa Cruz (Dallas, TX, USA) | Cat# sc-47778; RRID: AB_626632 |
| Luciferase (C-12) | Santa Cruz (Dallas, TX, USA) | Cat# sc-74548 |
| IgG (3E8) | Santa Cruz (Dallas, TX, USA) | Cat# sc-69786; RRID: AB_1124809 |
| IRDye 680RD | Li-cor (Lincoln, NE, USA) | Cat# 926-68071; RRID: AB_10956166 |
| IRDye 680LT | Li-cor (Lincoln, NE, USA) | Cat# 926-68022; RRID: AB_10715072 |
| IRDye 800CW | Li-cor (Lincoln, NE, USA) | Cat# 926-32211; RRID: AB_621843 |
| **Virus strains** |  |  |
| *HIV replication-competent virus (HIV strain 93/TH/051; R5- and X4-tropic virus isolated from a seropositive individual in Thailand)* | NIH AIDS reagent | ARP-2165 |
| **Chemicals, peptides, and recombinant protein** |  |  |
| Triton X-100 | Sigma-Aldrich (Burlington, MA, USA) | Cat# T928 |
| RPMI1640 | Invitrogen (Waltham, MA, USA) | Cat# 11835-030 |
| **DMEM** | Lonza (Walkersville, MD, USA) | Cat# 12–604F |
| **Opti-MEM (1X) +GlutaMAX™** | Gibco (Waltham, MA, USA) | Cat# 51985–034 |
| **HEPES Buffer** | Corning, New York, U.S. | Cat# 25-060-CI |
| **Fetal Bovine serum** | Gibco (Waltham, MA, USA) | Cat# 10082147 |
| **Pen strep** | Gibco (Waltham, MA, USA) | Cat# 15140–122 |
| **DMSO** | Sigma-Aldrich (Burlington, MA, USA) | Cat# D2650 |
| **BSA** | Sigma-Aldrich (Burlington, MA, USA) | Cat# A9647 |
| **2-Mercaptoethanol** | Sigma-Aldrich (Burlington, MA, USA) | Cat#M3148 |
| **Cocaine** | NIDA | N/A |
| **Nedisertib (M3814)** | Selleckchem (Houston, TX, USA) | Cat#S8586 |
| **NU7441 (KU-57788)** | Selleckchem (Houston, TX, USA) | Cat#S2638 |
| **Protein A-Sepharose 4B Conjugate** | Thermo Scientific (Waltham, MA, USA) | Cat# 101041 |
| **PageRuler™ Prestained Protein Ladder** | Thermo Scientific (Waltham, MA, USA) | Cat# 26617 |
| **Passive Lysis Buffer, 5X** | Promega (Madison, WI, USA) | Cat# E194A |
| **Trypan Blue stain** | Gibco (Waltham, MA, USA) | Cat# 15250–061 |
| **Lipofectamine 2000** | Thermo Scientific (Waltham, MA, USA) | Cat# 11668–019 |
| **Critical commercial assays** |  |  |
| **RNeasy mini kit** | Qiagen (Hilden, Germany) | Cat# 74104 |
| **RevertAid First Strand cDNA Synthesis Kit** | Thermo Scientific (Waltham, MA, USA) | Cat# K1622 |
| **BCA Protein Assay Kit** | Thermo Scientific (Waltham, MA, USA) | Cat# 23225 |
| **Luciferase Assay System** | Promega (Madison, WI, USA) | Cat# E1501 |
| **CellTiter 96® Aqueous One Solution Reagent** | Promega (Madison, WI, USA) | Cat# G5421 |
| **PowerUp SYBR Green Master Mix** | Thermo Scientific (Waltham, MA, USA) | Cat# 100029284 |
| **Experimental models: Cell lines** |  |  |
| **Jurkat-pHR'-P-Luc** | In-house generated model | N/A |
| **2D10 cells line** | In-house generated model | N/A |
| **Jurkat cells line** | ATCC | TIB-152 |
| **MT-4 cell line** | NIH AIDS reagent | ARP-120 |
| **THP-1 cell line** | ATCC | TIB-202 |
| **Microglial cells** | ATCC |  |
| **PBMCs** | Lonza (Cambridge, MA) | Cat# CC-2702 |
| **Software and algorithms** |  |  |
| **Prism 9** | Graphpad | Ver 9.1.2 (226) |
| **Odyssey infrared imaging software** | Li-cor | Ver 3.0.30 |
| **ImageJ software** | NIH | 1.53e |

## Notes

### Competing Interest Statement

The authors have declared no competing interest.

## References

1. Tyagi M, Bukrinsky M: Human immunodeficiency virus (HIV) latency: the major hurdle in HIV eradication. Mol Med 2012, 18(1):1096–1108.

2. Mbonye U, Karn J: Control of HIV latency by epigenetic and non-epigenetic mechanisms. Curr HIV Res 2011, 9(8):554–567.

3. Nguyen K, Dobrowolski C, Shukla M, Cho WK, Luttge B, Karn J: Inhibition of the H3K27 demethylase UTX enhances the epigenetic silencing of HIV proviruses and induces HIV-1 DNA hypermethylation but fails to permanently block HIV reactivation. PLoS Pathog 2021, 17(10):e1010014.

4. Choudhary SK, Margolis DM: Curing HIV: Pharmacologic approaches to target HIV-1 latency. Annu Rev Pharmacol Toxicol 2011, 51:397–418.

5. Hokello J, Sharma AL, Tyagi M: An Update on the HIV DNA Vaccine Strategy. Vaccines (Basel) 2021, 9(6).

6. Walensky RP, Paltiel AD, Losina E, Mercincavage LM, Schackman BR, Sax PE, Weinstein MC, Freedberg KA: The survival benefits of AIDS treatment in the United States. J Infect Dis 2006, 194(1):11–19.

7. Saag MS: HIV 101: fundamentals of antiretroviral therapy. Top Antivir Med 2019, 27(3):123- 127.

8. Wilson EM, Sereti I: Immune restoration after antiretroviral therapy: the pitfalls of hasty or incomplete repairs. Immunol Rev 2013, 254(1):343–354.

9. Gay CL, Cohen MS: Antiretrovirals to prevent HIV infection: pre- and postexposure prophylaxis. Curr Infect Dis Rep 2008, 10(4):323–331.

10. Bertoni N, Singer M, Silva CM, Clair S, Malta M, Bastos FI: Knowledge of AIDS and HIV transmission among drug users in Rio de Janeiro, Brazil. Harm Reduct J 2011, 8:5.

11. Iskandar S, Basar D, Hidayat T, Siregar IM, Pinxten L, van Crevel R, Van der Ven AJ, De Jong CA: High risk behavior for HIV transmission among former injecting drug users: a survey from Indonesia. BMC Public Health 2010, 10:472.

12. Baggaley RF, Boily MC, White RG, Alary M: Risk of HIV-1 transmission for parenteral exposure and blood transfusion: a systematic review and meta-analysis. AIDS 2006, 20(6):805–812.

13. Hudgins R, McCusker J, Stoddard A: Cocaine use and risky injection and sexual behaviors. Drug Alcohol Depend 1995, 37(1):7–14.

14. Purcell DW, Parsons JT, Halkitis PN, Mizuno Y, Woods WJ: Substance use and sexual transmission risk behavior of HIV-positive men who have sex with men. J Subst Abuse 2001, 13(1-2):185–200.

15. Molitor F, Truax SR, Ruiz JD, Sun RK: Association of methamphetamine use during sex with risky sexual behaviors and HIV infection among non-injection drug users. West J Med 1998, 168(2):93–97.

16. Tyagi M, Weber J, Bukrinsky M, Simon GL: The effects of cocaine on HIV transcription. J Neurovirol 2016, 22(3):261–274.

17. Tyagi M, Bukrinsky M, Simon GL: Mechanisms of HIV Transcriptional Regulation by Drugs of Abuse. Curr HIV Res 2016, 14(5):442–454.

18. Ciccarone D: Stimulant abuse: pharmacology, cocaine, methamphetamine, treatment, attempts at pharmacotherapy. Prim Care 2011, 38(1):41-58.

19. Pomara C, Cassano T, D’Errico S, Bello S, Romano AD, Riezzo I, Serviddio G: Data available on the extent of cocaine use and dependence: biochemistry, pharmacologic effects and global burden of disease of cocaine abusers. Curr Med Chem 2012, 19(33):5647–5657.

20. Calatayud J, Gonzalez A: History of the development and evolution of local anesthesia since the coca leaf. Anesthesiology 2003, 98(6):1503–1508.

21. Goldstein RA, DesLauriers C, Burda AM: Cocaine: history, social implications, and toxicity--a review. Dis Mon 2009, 55(1):6–38.

22. Jeffcoat AR, Perez-Reyes M, Hill JM, Sadler BM, Cook CE: Cocaine disposition in humans after intravenous injection, nasal insufflation (snorting), or smoking. Drug Metab Dispos 1989, 17(2):153–159.

23. Klinkenberg WD, Sacks S, Hiv/Aids Treatment Adherence HO, Cost Study G: Mental disorders and drug abuse in persons living with HIV/AIDS. AIDS Care 2004, 16 Suppl 1:S22–42.

24. Khalsa JH, Elkashef A: Interventions for HIV and hepatitis C virus infections in recreational drug users. Clin Infect Dis 2010, 50(11):1505–1511.

25. Buch S, Yao H, Guo M, Mori T, Mathias-Costa B, Singh V, Seth P, Wang J, Su TP: Cocaine and HIV-1 interplay in CNS: cellular and molecular mechanisms. Curr HIV Res 2012, 10(5):425–428.

26. Parikh N, Nonnemacher MR, Pirrone V, Block T, Mehta A, Wigdahl B: Substance abuse, HIV-1 and hepatitis. Curr HIV Res 2012, 10(7):557-571.

27. Sharma AL, Shafer D, Netting D, Tyagi M: Cocaine sensitizes the CD4(+) T cells for HIV infection by co-stimulating NFAT and AP-1. iScience 2022, 25(12):105651.

28. Sahu G, Farley K, El-Hage N, Aiamkitsumrit B, Fassnacht R, Kashanchi F, Ochem A, Simon GL, Karn J, Hauser KF et al: Cocaine promotes both initiation and elongation phase of HIV-1 transcription by activating NF-kappaB and MSK1 and inducing selective epigenetic modifications at HIV-1 LTR. Virology 2015, 483:185–202.

29. Dash S, Balasubramaniam M, Villalta F, Dash C, Pandhare J: Impact of cocaine abuse on HIV pathogenesis. Front Microbiol 2015, 6:1111.

30. Sonti S, Tyagi K, Pande A, Daniel R, Sharma AL, Tyagi M: Crossroads of Drug Abuse and HIV Infection: Neurotoxicity and CNS Reservoir. Vaccines (Basel) 2022, 10(2).

31. Riedl T, Egly JM: Phosphorylation in transcription: the CTD and more. Gene Expr 2000, 9(1- 2):3–13.

32. Brookes E, Pombo A: Modifications of RNA polymerase II are pivotal in regulating gene expression states. EMBO Rep 2009, 10(11):1213–1219.

33. Hsin JP, Manley JL: The RNA polymerase II CTD coordinates transcription and RNA processing. Genes Dev 2012, 26(19):2119–2137.

34. Dahmus ME: Phosphorylation of the C-terminal domain of RNA polymerase II. Biochim Biophys Acta 1995, 1261(2):171-182.

35. Orphanides G, Lagrange T, Reinberg D: The general transcription factors of RNA polymerase II. Genes Dev 1996, 10(21):2657–2683.

36. Isel C, Karn J: Direct evidence that HIV-1 Tat stimulates RNA polymerase II carboxyl-terminal domain hyperphosphorylation during transcriptional elongation. J Mol Biol 1999, 290(5):929–941.

37. Tyagi S, Ochem A, Tyagi M: DNA-dependent protein kinase interacts functionally with the RNA polymerase II complex recruited at the human immunodeficiency virus (HIV) long terminal repeat and plays an important role in HIV gene expression. J Gen Virol 2011, 92(Pt 7):1710–1720.

38. Nagasawa M, Watanabe F, Suwa A, Yamamoto K, Tsukada K, Teraoka H: Nuclear translocation of the catalytic component of DNA-dependent protein kinase upon growth stimulation in normal human T lymphocytes. Cell Struct Funct 1997, 22(6):585–594.

39. Zicari S, Sharma AL, Sahu G, Dubrovsky L, Sun L, Yue H, Jada T, Ochem A, Simon G, Bukrinsky M et al: DNA dependent protein kinase (DNA-PK) enhances HIV transcription by promoting RNA polymerase II activity and recruitment of transcription machinery at HIV LTR. Oncotarget 2020, 11(7):699–726.

40. Hartley KO, Gell D, Smith GC, Zhang H, Divecha N, Connelly MA, Admon A, Lees-Miller SP, Anderson CW, Jackson SP: DNA-dependent protein kinase catalytic subunit: a relative of phosphatidylinositol 3-kinase and the ataxia telangiectasia gene product. Cell 1995, 82(5):849–856.

41. Gottlieb TM, Jackson SP: The DNA-dependent protein kinase: requirement for DNA ends and association with Ku antigen. Cell 1993, 72(1):131–142.

42. Reeves WH, Sthoeger ZM: Molecular cloning of cDNA encoding the p70 (Ku) lupus autoantigen. J Biol Chem 1989, 264(9):5047–5052.

43. Lees-Miller SP, Godbout R, Chan DW, Weinfeld M, Day RS, 3rd, Barron GM, Allalunis-Turner J: Absence of p350 subunit of DNA-activated protein kinase from a radiosensitive human cell line. Science 1995, 267(5201):1183-1185.

44. Taccioli GE, Gottlieb TM, Blunt T, Priestley A, Demengeot J, Mizuta R, Lehmann AR, Alt FW, Jackson SP, Jeggo PA: Ku80: product of the XRCC5 gene and its role in DNA repair and V(D)J recombination. Science 1994, 265(5177):1442-1445.

45. Blackford AN, Jackson SP: ATM, ATR, and DNA-PK: The Trinity at the Heart of the DNA Damage Response. Mol Cell 2017, 66(6):801–817.

46. Dylgjeri E, Knudsen KE: DNA-PKcs: A Targetable Protumorigenic Protein Kinase. Cancer Res 2022, 82(4):523–533.

47. Dvir A, Stein LY, Calore BL, Dynan WS: Purification and characterization of a template- associated protein kinase that phosphorylates RNA polymerase II. J Biol Chem 1993, 268(14):10440–10447.

48. Peterlin BM, Price DH: Controlling the elongation phase of transcription with P-TEFb. Mol Cell 2006, 23(3):297–305.

49. Kim YK, Bourgeois CF, Isel C, Churcher MJ, Karn J: Phosphorylation of the RNA polymerase II carboxyl-terminal domain by CDK9 is directly responsible for human immunodeficiency virus type 1 Tat-activated transcriptional elongation. Mol Cell Biol 2002, 22(13):4622–4637.

50. Yamamoto S, Watanabe Y, van der Spek PJ, Watanabe T, Fujimoto H, Hanaoka F, Ohkuma Y: Studies of nematode TFIIE function reveal a link between Ser-5 phosphorylation of RNA polymerase II and the transition from transcription initiation to elongation. Mol Cell Biol 2001, 21(1):1–15.

51. Kim YK, Bourgeois CF, Pearson R, Tyagi M, West MJ, Wong J, Wu SY, Chiang CM, Karn J: Recruitment of TFIIH to the HIV LTR is a rate-limiting step in the emergence of HIV from latency. EMBO J 2006, 25(15):3596–3604.

52. Saunders A, Core LJ, Lis JT: Breaking barriers to transcription elongation. Nat Rev Mol Cell Biol 2006, 7(8):557–567.

53. Kao SY, Calman AF, Luciw PA, Peterlin BM: Anti-termination of transcription within the long terminal repeat of HIV-1 by tat gene product. Nature 1987, 330(6147):489-493.

54. Guenther MG, Levine SS, Boyer LA, Jaenisch R, Young RA: A chromatin landmark and transcription initiation at most promoters in human cells. Cell 2007, 130(1):77–88.

55. Muse GW, Gilchrist DA, Nechaev S, Shah R, Parker JS, Grissom SF, Zeitlinger J, Adelman K: RNA polymerase is poised for activation across the genome. Nat Genet 2007, 39(12):1507–1511.

56. Wu CH, Yamaguchi Y, Benjamin LR, Horvat-Gordon M, Washinsky J, Enerly E, Larsson J, Lambertsson A, Handa H, Gilmour D: NELF and DSIF cause promoter proximal pausing on the hsp70 promoter in Drosophila. Genes Dev 2003, 17(11):1402–1414.

57. Yamaguchi Y, Shibata H, Handa H: Transcription elongation factors DSIF and NELF: promoter- proximal pausing and beyond. Biochim Biophys Acta 2013, 1829(1):98–104.

58. Bunch H, Calderwood SK: TRIM28 as a novel transcriptional elongation factor. BMC Mol Biol 2015, 16:14.

59. Bunch H, Lawney BP, Lin YF, Asaithamby A, Murshid A, Wang YE, Chen BP, Calderwood SK: Transcriptional elongation requires DNA break-induced signalling. Nat Commun 2015, 6:10191.

60. Bunch H, Zheng X, Burkholder A, Dillon ST, Motola S, Birrane G, Ebmeier CC, Levine S, Fargo D, Hu G et al: TRIM28 regulates RNA polymerase II promoter-proximal pausing and pause release. Nat Struct Mol Biol 2014, 21(10):876–883.

61. Czerwinska P, Mazurek S, Wiznerowicz M: The complexity of TRIM28 contribution to cancer. J Biomed Sci 2017, 24(1):63.

62. Allouch A, Di Primio C, Alpi E, Lusic M, Arosio D, Giacca M, Cereseto A: The TRIM family protein KAP1 inhibits HIV-1 integration. Cell Host Microbe 2011, 9(6):484–495.

63. Randolph K, Hyder U, D’Orso I: KAP1/TRIM28: Transcriptional Activator and/or Repressor of Viral and Cellular Programs? Front Cell Infect Microbiol 2022, 12:834636.

64. Ait-Ammar A, Bellefroid M, Daouad F, Martinelli V, Van Assche J, Wallet C, Rodari A, De Rovere M, Fahrenkrog B, Schwartz C et al: Inhibition of HIV-1 gene transcription by KAP1 in myeloid lineage. Sci Rep 2021, 11(1):2692.

65. Parada CA, Roeder RG: Enhanced processivity of RNA polymerase II triggered by Tat-induced phosphorylation of its carboxy-terminal domain. Nature 1996, 384(6607):375-378.

66. Ivanov D, Kwak YT, Guo J, Gaynor RB: Domains in the SPT5 protein that modulate its transcriptional regulatory properties. Mol Cell Biol 2000, 20(9):2970–2983.

67. Fujinaga K, Irwin D, Huang Y, Taube R, Kurosu T, Peterlin BM: Dynamics of human immunodeficiency virus transcription: P-TEFb phosphorylates RD and dissociates negative effectors from the transactivation response element. Mol Cell Biol 2004, 24(2):787–795.

68. Tyagi M, Pearson RJ, Karn J: Establishment of HIV latency in primary CD4+ cells is due to epigenetic transcriptional silencing and P-TEFb restriction. J Virol 2010, 84(13):6425–6437.

69. Sobhian B, Laguette N, Yatim A, Nakamura M, Levy Y, Kiernan R, Benkirane M: HIV-1 Tat assembles a multifunctional transcription elongation complex and stably associates with the 7SK snRNP. Mol Cell 2010, 38(3):439–451.

70. He N, Chan CK, Sobhian B, Chou S, Xue Y, Liu M, Alber T, Benkirane M, Zhou Q: Human Polymerase-Associated Factor complex (PAFc) connects the Super Elongation Complex (SEC) to RNA polymerase II on chromatin. Proc Natl Acad Sci U S A 2011, 108(36):E636–645.

71. Chou S, Upton H, Bao K, Schulze-Gahmen U, Samelson AJ, He N, Nowak A, Lu H, Krogan NJ, Zhou Q et al: HIV-1 Tat recruits transcription elongation factors dispersed along a flexible AFF4 scaffold. Proc Natl Acad Sci U S A 2013, 110(2):E123–131.

72. Tyagi M, Rusnati M, Presta M, Giacca M: Internalization of HIV-1 tat requires cell surface heparan sulfate proteoglycans. J Biol Chem 2001, 276(5):3254–3261.

73. Barboric M, Peterlin BM: A new paradigm in eukaryotic biology: HIV Tat and the control of transcriptional elongation. PLoS Biol 2005, 3(2):e76.

74. Tahirov TH, Babayeva ND, Varzavand K, Cooper JJ, Sedore SC, Price DH: Crystal structure of HIV-1 Tat complexed with human P-TEFb. Nature 2010, 465(7299):747-751.

75. Karn J: Tackling Tat. J Mol Biol 1999, 293(2):235-254.

76. Taube R, Fujinaga K, Wimmer J, Barboric M, Peterlin BM: Tat transactivation: a model for the regulation of eukaryotic transcriptional elongation. Virology 1999, 264(2):245–253.

77. Dull T, Zufferey R, Kelly M, Mandel RJ, Nguyen M, Trono D, Naldini L: A third-generation lentivirus vector with a conditional packaging system. J Virol 1998, 72(11):8463–8471.

78. Tyagi M, Karn J: CBF-1 promotes transcriptional silencing during the establishment of HIV-1 latency. EMBO J 2007, 26(24):4985–4995.

79. Hokello J, Lakhikumar Sharma A, Tyagi M: AP-1 and NF-kappaB synergize to transcriptionally activate latent HIV upon T-cell receptor activation. FEBS Lett 2021, 595(5):577–594.

80. Pearson R, Kim YK, Hokello J, Lassen K, Friedman J, Tyagi M, Karn J: Epigenetic silencing of human immunodeficiency virus (HIV) transcription by formation of restrictive chromatin structures at the viral long terminal repeat drives the progressive entry of HIV into latency. J Virol 2008, 82(24):12291–12303.

81. Tyagi M, Karn J: CBF-1 promotes transcriptional silencing during the establishment of HIV-1 latency. EMBO J 2007, 26(24):4985–4995.

82. Carr MI, Zimmermann A, Chiu LY, Zenke FT, Blaukat A, Vassilev LT: DNA-PK Inhibitor, M3814, as a New Combination Partner of Mylotarg in the Treatment of Acute Myeloid Leukemia. Front Oncol 2020, 10:127.

83. Sun Q, Guo Y, Liu X, Czauderna F, Carr MI, Zenke FT, Blaukat A, Vassilev LT: Therapeutic Implications of p53 Status on Cancer Cell Fate Following Exposure to Ionizing Radiation and the DNA-PK Inhibitor M3814. Mol Cancer Res 2019, 17(12):2457–2468.

84. van Bussel MTJ, Awada A, de Jonge MJA, Mau-Sorensen M, Nielsen D, Schoffski P, Verheul HMW, Sarholz B, Berghoff K, El Bawab S et al: A first-in-man phase 1 study of the DNA-dependent protein kinase inhibitor peposertib (formerly M3814) in patients with advanced solid tumours. Br J Cancer 2021, 124(4):728–735.

85. Wise HC, Iyer GV, Moore K, Temkin SM, Gordon S, Aghajanian C, Grisham RN: Activity of M3814, an Oral DNA-PK Inhibitor, In Combination with Topoisomerase II Inhibitors in Ovarian Cancer Models. Sci Rep 2019, 9(1):18882.

86. Zenke FT, Zimmermann A, Sirrenberg C, Dahmen H, Kirkin V, Pehl U, Grombacher T, Wilm C, Fuchss T, Amendt C et al: Pharmacologic Inhibitor of DNA-PK, M3814, Potentiates Radiotherapy and Regresses Human Tumors in Mouse Models. Mol Cancer Ther 2020, 19(5):1091-1101.

87. Hardcastle IR, Cockcroft X, Curtin NJ, El-Murr MD, Leahy JJJ, Stockley M, Golding BT, Rigoreau L, Richardson C, Smith GCM et al: Discovery of potent chromen-4-one inhibitors of the DNA-dependent protein kinase (DNA-PK) using a small-molecule library approach. Journal of Medicinal Chemistry 2005, 48(24):7829–7846.

88. Zhao Y, Thomas HD, Batey MA, Cowell IG, Richardson CJ, Griffin RJ, Calvert AH, Newell DR, Smith GC, Curtin NJ: Preclinical evaluation of a potent novel DNA-dependent protein kinase inhibitor NU7441. Cancer Res 2006, 66(10):5354–5362.

89. Davidson D, Amrein L, Panasci L, Aloyz R: Small Molecules, Inhibitors of DNA-PK, Targeting DNA Repair, and Beyond. Front Pharmacol 2013, 4:5.

90. Cornell L, Munck J, Curtin N, Reeves H: DNA-Pk or Atm Inhibition Inhibits Non-Homologous End Joining and Enhances Chemo- and Radio Sensitivity in Hepatocellular Cancer Cell Lines. Gut 2012, 61:A201–A201.

91. Feng FY, Brenner JC, Hussain M, Chinnaiyan AM: Molecular pathways: targeting ETS gene fusions in cancer. Clinical cancer research : an official journal of the American Association for Cancer Research 2014, 20(17):4442–4448.

92. Jekimovs C, Bolderson E, Suraweera A, Adams M, O’Byrne KJ, Richard DJ: Chemotherapeutic compounds targeting the DNA double-strand break repair pathways: the good, the bad, and the promising. Front Oncol 2014, 4:86.

93. Sharma AL, Hokello J, Sonti S, Zicari S, Sun L, Alqatawni A, Bukrinsky M, Simon G, Chauhan A, Daniel R et al: CBF-1 Promotes the Establishment and Maintenance of HIV Latency by Recruiting Polycomb Repressive Complexes, PRC1 and PRC2, at HIV LTR. Viruses 2020, 12(9).

94. Hokello J, Tyagi P, Dimri S, Sharma AL, Tyagi M: Comparison of the Biological Basis for Non-HIV Transmission to HIV-Exposed Seronegative Individuals, Disease Non-Progression in HIV Long-Term Non-Progressors and Elite Controllers. Viruses 2023, 15(6).

95. Sharma AL, Hokello J, Tyagi M: Circumcision as an Intervening Strategy against HIV Acquisition in the Male Genital Tract. Pathogens 2021, 10(7).

96. Hokello J, Sharma AL, Dimri M, Tyagi M: Insights into the HIV Latency and the Role of Cytokines. Pathogens 2019, 8(3).

97. Sharma AL, Singh TR, Singh LS: Antiretroviral resistance, genotypic characterization and origin of Human Immunodeficiency Virus among the infected wives of Intravenous drug users in Manipur. Sci Rep 2018, 8(1):15183.

98. Hokello J, Sharma AL, Tyagi M: Efficient Non-Epigenetic Activation of HIV Latency through the T-Cell Receptor Signalosome. Viruses 2020, 12(8).

99. Liu H, Miller LG, Hays RD, Golin CE, Wu T, Wenger NS, Kaplan AH: Repeated measures longitudinal analyses of HIV virologic response as a function of percent adherence, dose timing, genotypic sensitivity, and other factors. J Acquir Immune Defic Syndr 2006, 41(3):315–322.

100. Parruti G, Manzoli L, Toro PM, D’Amico G, Rotolo S, Graziani V, Schioppa F, Consorte A, Alterio L, Toro GM et al: Long-term adherence to first-line highly active antiretroviral therapy in a hospital-based cohort: predictors and impact on virologic response and relapse. AIDS Patient Care STDS 2006, 20(1):48–56.

101. Jackson SP, MacDonald JJ, Lees-Miller S, Tjian R: GC box binding induces phosphorylation of Sp1 by a DNA-dependent protein kinase. Cell 1990, 63(1):155–165.

102. Goodwin JF, Knudsen KE: Beyond DNA repair: DNA-PK function in cancer. Cancer Discov 2014, 4(10):1126–1139.

103. Marshall NF, Peng J, Xie Z, Price DH: Control of RNA polymerase II elongation potential by a novel carboxyl-terminal domain kinase. J Biol Chem 1996, 271(43):27176–27183.

104. Marshall NF, Price DH: Purification of P-TEFb, a transcription factor required for the transition into productive elongation. J Biol Chem 1995, 270(21):12335–12338.

