## Supplementary material for "Cocaine-induced DNA-PK relieves RNAP II pausing by promoting TRIM28 phosphorylation": Sharma et. al Supplementary file

Research Article

Supplementary Table S1

**Table S1: List of primer sequences.**

| Primer Sets | Primer name | Sequence (5'-3') | Purpose |
|---|---|---|---|
| **1st Set primer** | GAPDHF | CGGGATTGTCTGCCCTAATTAT | Real time PCR |
| | GAPDHR | GCACGGAAGGTCACGATGT | Real time PCR |
| **2nd Set primer** | HIV Promoter F | AGCTTGCTACAAGGGACTTTCC | Real time PCR |
| | HIV promoter R | ACCCAGTACAGGCAAAAAGCAG | Real time PCR |
| **3rd Set primer** | HIV Nuc-1F | CTGGGAGCTCTCTGGCTAACTA | Real time PCR |
| | HIV Nuc-1R | TTACCAGAGTCACACAACAGACG | Real time PCR |
| **4th Set primer** | HIV Nuc-2F | GACTGGTGAGTACGCCAAAA | Real time PCR |
| | HIV Nuc-2R | TTTCCCACTGCGATCTAATTC | Real time PCR |
| **5th Set primer** | HIV envF | TGAGGGACAATCGGAGAAG | Real time PCR |
| | HIV envR | TCTGCACCACTCTTCTCTT | Real time PCR |
| **6th Set primer** | DNA-PKF1 | ACGGTAGGGGAAAGCCATTG | Real time PCR |
| | DNA-PKR1 | CGCTATAGGTCCTCAGCTGC | Real time PCR |
| **7th Set primer** | ActinF1 | AGAGCAAGAGAGGCATCCTG | Real time PCR |
| | ActinR1 | GGGTCATCTTTTCACGGTTGG | Real time PCR |

**Supplementary Figure S1**

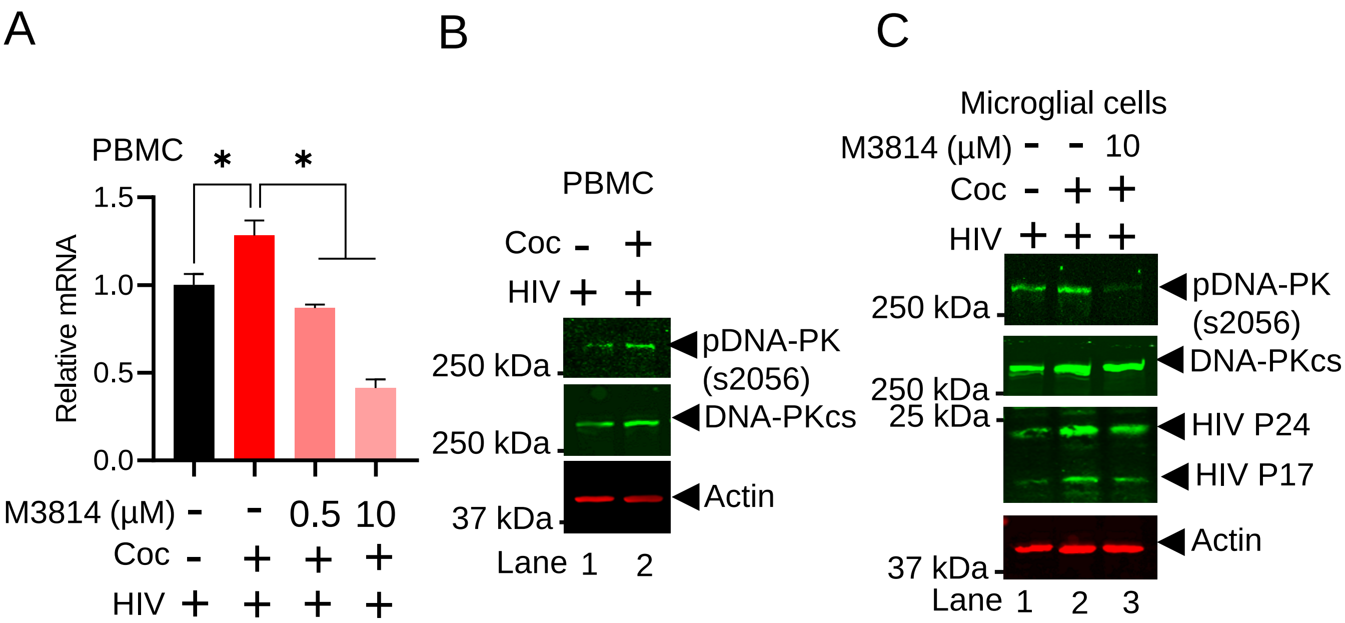


**Supplementary Figure S1**: **DNA-PK inhibition strongly suppresses cocaine induced HIV transcription and replication**.

(**A**) PBMCs were treated with M3814 for overnight 24 h. Next day cells were treated with cocaine for 3 h and thereafter infected with replication competent HIV. DNA-PK (**A**) and HIV transcripts (**in main figure 5D and E**) were quantified by real-time PCR. The result is expressed as mean ± SD and analyzed by two-way ANOVA followed by Tukey's multiple comparisons test. Asterisks over the bars indicate significant differences: $*p < 0.05$. (**B**) PBMCs were exposed to cocaine for a duration of 3 h, followed by infection with HIV. Subsequently, nuclear lysates were subjected to analysis via immunoblotting using specific antibodies targeting phosphorylated pDNA-PK (S2056), DNA-PKcs, and Actin. (**C**) Microglial cells were exposed to M3814 for a duration of 24 hours. The following day, the cells were subjected to treatment with cocaine and subsequent infection with HIV. Nuclear lysates were then analyzed via immunoblotting, utilizing specific antibodies targeting phosphorylated pDNA-PK (S2056), DNA-PKcs, HIV p24, and Actin.
